# IPMK physically binds to the SWI/SNF complex and modulates BRG1 occupancy

**DOI:** 10.1101/2021.09.15.460446

**Authors:** Jiyoon Beon, Sungwook Han, Seung Eun Park, Kwangbeom Hyun, Song-Yi Lee, Hyun-Woo Rhee, Jeong Kon Seo, Jaehoon Kim, Seyun Kim, Daeyoup Lee

## Abstract

Inositol polyphosphate multikinase (IPMK), a key enzyme in the inositol polyphosphate (IP) metabolism, is a pleiotropic signaling factor involved in major biological events including transcriptional control. In yeasts, IPMK and its IP products were known to promote the activity of SWI/SNF chromatin remodeling complex, which plays a critical role in gene expression by regulating chromatin accessibility. However, the direct linkage between IPMK and chromatin remodelers remains unclear, raising a question on how IPMK contributes to the transcriptional regulation in mammals. By employing unbiased screenings and in vivo/in vitro immunoprecipitations, here we demonstrated that IPMK physically associates with native mammalian SWI/SNF complexes by directly binding to SMARCB1, BRG1, and SMARCC1. Furthermore, we identified the specific domains required for the IPMK-SMARCB1 binding. Notably, using CUT&RUN and ATAC-seq assays, we discovered that IPMK co-localizes with BRG1 and regulates BRG1 localization as well as BRG1-mediated chromatin accessibility in a genome-wide manner (including promoter-TSS) in mouse embryonic stem cells. Finally, our mRNA-seq analyses revealed that IPMK and SMARCB1 regulate common gene sets, validating a functional link between IPMK and SWI/SNF complex. Together, these findings establish an importance of IPMK in promoter targeting of the SWI/SNF complex, thereby contributing to SWI/SNF-meditated chromatin accessibility and transcription.

## INTRODUCTION

Inositol polyphosphates are a class of signaling messengers that mediate diverse biological events such as cellular growth, proliferation, and metabolic homeostasis. Inositol polyphosphate multikinase (IPMK) is an essential enzyme for the synthesis of these inositol polyphosphates including inositol tetrakisphosphates (IP_4_, both Ins(1,3,4,5)P­and Ins(1,4,5,6)P­) and pentakisphosphates (IP_5_, Ins(1,3,4,5,6)P_5_) (Chakraborty, Kim, & Snyder, 2011; Hatch & York, 2010; Saiardi, Erdjument-Bromage, Snowman, Tempst, & Snyder, 1999). In addition to its role as a phosphatidylinositol 3-kinase (thereby producing phosphatidylinositol 3,4,5-trisphosphate (PIP_3_)), IPMK also non-catalytically controls various signaling factors including the mammalian target of rapamycin (mTOR), AMP-activated protein kinase (AMPK), and TRAF6 (Bang et al., 2012; E. Kim, Ahn, Kim, Lee, & Kim, 2017; E. Kim, Beon, et al., 2017; S. Kim et al., 2011; Maag et al., 2011; Resnick et al., 2005). These findings suggest that IPMK plays a critical role in coordinating the major biological events.

Increasing evidence strongly indicates that nuclear IPMK act as a key factor in gene expression regulation. IPMK was originally cloned from yeast as Arg82 (yeast IPMK), a gene required for the regulation of arginine metabolism (Bechet, Greenson, & Wiame, 1970; Dubois, Bercy, & Messenguy, 1987; Odom, Stahlberg, Wente, & York, 2000). The physical interaction between Arg82 and the yeast transcription factor MCM1, a yeast homolog of the mammalian serum response factor (SRF), is crucial for the transcriptional control (Bercy, Dubois, & Messenguy, 1987; Christ & Tye, 1991; Messenguy & Dubois, 1993; Odom et al., 2000). In mammals, IPMK-SRF binding was found to be a critical event for SRF-dependent gene induction (E. Kim et al., 2013). Furthermore, other functions of nuclear IPMK are mediated by its diverse interactions with p53, steroidogenic factor 1, and CBP/p300 (Blind, 2014; Blind et al., 2014; Malabanan & Blind, 2016; Xu, Paul, et al., 2013; Xu, Sen, et al., 2013; Xu & Snyder, 2013).

Chromatin remodeling is essential for the efficient transcription of eukaryotic genes (Kouzarides, 2007; Trotter & Archer, 2007; Vignali, Hassan, Neely, & Workman, 2000). Particularly, SWI/SNF is a large family of ATP-dependent chromatin remodeling complexes that have been characterized as transcriptional activators or repressors. These complexes enable the transcription machinery or other transcription factors to gain access to their target genes (Arnaud, Le Loarer, & Tirode, 2018; Hargreaves & Crabtree, 2011). In mammalian cells, the canonical SWI/SNF complex contains one of the two mutually exclusive ATPases, BRM (SMARCA2) or BRG1 (SMARCA4), in addition to a core set of subunits consisting of BAF155 (BRG1-associated factor or SMARCC1), SMARCB1 (hSNF5 or INI1), and BAF170 (SMARCC2), as well as four to eight other accessory subunits (Khavari, Peterson, Tamkun, Mendel, & Crabtree, 1993; W. Wang, Côté, et al., 1996; W. Wang, Xue, et al., 1996). Importantly, this SWI/SNF complex mediates nucleosome structure modifications and regulates the positioning of nucleosomes in an ATP-dependent manner, thereby modulating the accessibility of regulatory proteins. Therefore, the SWI/SNF chromatin remodeling complex is critical for various biological processes, including gene transcription, cell cycle regulation, and cell differentiation (Ho, Jothi, et al., 2009; Ho, Ronan, et al., 2009; Hodges, Kirkland, & Crabtree, 2016; K. H. Kim & Roberts, 2014; Tolstorukov et al., 2013; X. Wang, Haswell, & Roberts, 2014).

Despite the importance of both inositol polyphosphates and chromatin remodeling in transcriptional regulation, only a few studies have addressed the linkage between inositol polyphosphates and chromatin remodeling. Previous study in yeasts demonstrated that inositol polyphosphates could regulate the nucleosome-sliding activity of chromatin remodeling complexes *in vitro* (Shen, Xiao, Ranallo, Wu, & Wu, 2003). Specifically, inositol tetrakisphosphates and pentakisphosphates (IP_4_ and IP_5_) stimulate the activity of the SWI/SNF complex, whereas inositol hexakisphosphate (IP_6_) inhibits the activity of NURF, ISW2, and INO80 complexes. Another study in yeast illustrated that Arg82 (i.e., a yeast IPMK) mutation which led to defective IP_4_ and IP_5_ production, causes inefficient recruitment of the SWI/SNF complex, resulting in impaired chromatin remodeling at the phosphate-responsive *PHO5* gene promoter (Steger, Haswell, Miller, Wente, & O’Shea, 2003). In mammals, inositol hexakisphosphate kinase 1 (IP6K1) was recently found to directly interact with Jumonji domain containing 2C (JMJD2C), a histone demethylase. IP6K1 and its product, 5-IP_7_, appear to mediate JMJDC2-target gene expression in mammalian cells via regulating the chromatin association of JMJDC2 and levels of trimethyl-histone H3 lysine 9 (Burton, Azevedo, Andreassi, Riccio, & Saiardi, 2013). Taken together, these findings suggest that inositol polyphosphates and their enzymes play an important role in chromatin remodeling and transcription. However, direct linkages between IPMK (which produces inositol polyphosphate) and the chromatin remodeling complex SWI/SNF have not been previously reported, and it is still unclear whether IPMK contributes the transcriptional regulation of mammals.

To address these issues, we first performed unbiased screening assays and elucidated that the core subunits of the SWI/SNF complex, including SMARCB1 and BRG1, physically interact with IPMK. The physical association between IPMK and SWI/SNF complex was confirmed by our *in vitro* and *in vivo* immunoprecipitation assays. Furthermore, the specific binding sites between IPMK and SMARCB1 were mapped in detail. To investigate the biological role of IPMK-SWI/SNF complex binding, we performed various next-generation sequencing. We detected that IPMK and BRG1 were co-localized at the chromatin, especially at the promoter-TSS. Surprisingly, we found that IPMK depletion significantly reduced the global BRG1 occupancy and BRG1-mediated chromatin accessibility, especially at the bivalent promoters. The IPMK depletion also affected the transcription of genes with the reduced BRG1 occupancy and chromatin accessibility at their promoter-TSS. Lastly, we identified that IPMK and SMARCB1 regulate the common set of genes in the same manner. Taken together, our findings demonstrate the direct linkage between IPMK and the SWI/SNF complex (both physical and functional interactions), as well as the crucial role of IPMK in regulating BRG1 occupancy, BRG1-associated chromatin accessibility, and transcription.

## RESULTS

### Identification of IPMK-binding/interacting proteins

To investigate putative IPMK targets, we performed yeast two-hybrid screenings using IPMK as a bait and a human brain cDNA library as prey. The co-transformants of GAL4-DB fusion plasmid pGBKT7-IPMK (prey) and GAL4-AD fusion plasmid pACT2-SMARCB1 (bait) resulted in reporter gene activation, thus demonstrating cell growth on selective mediums, whereas co-transformants of pGBKT-7 and pACT2-SMARCB1 did not grow (Figure 1A). Approximately 23-36 proteins were identified as potential targets interacting with IPMK (Supplementary Table 1). Among these putative targets, only SMARCB1 was present in both duplicates of the yeast two-hybrid screening (Supplementary Table 1). These results indicate SMARCB1, a core subunit of SWI/SNF chromatin remodeler, as a potential candidate for IPMK-binding proteins.

**Figure 1.**
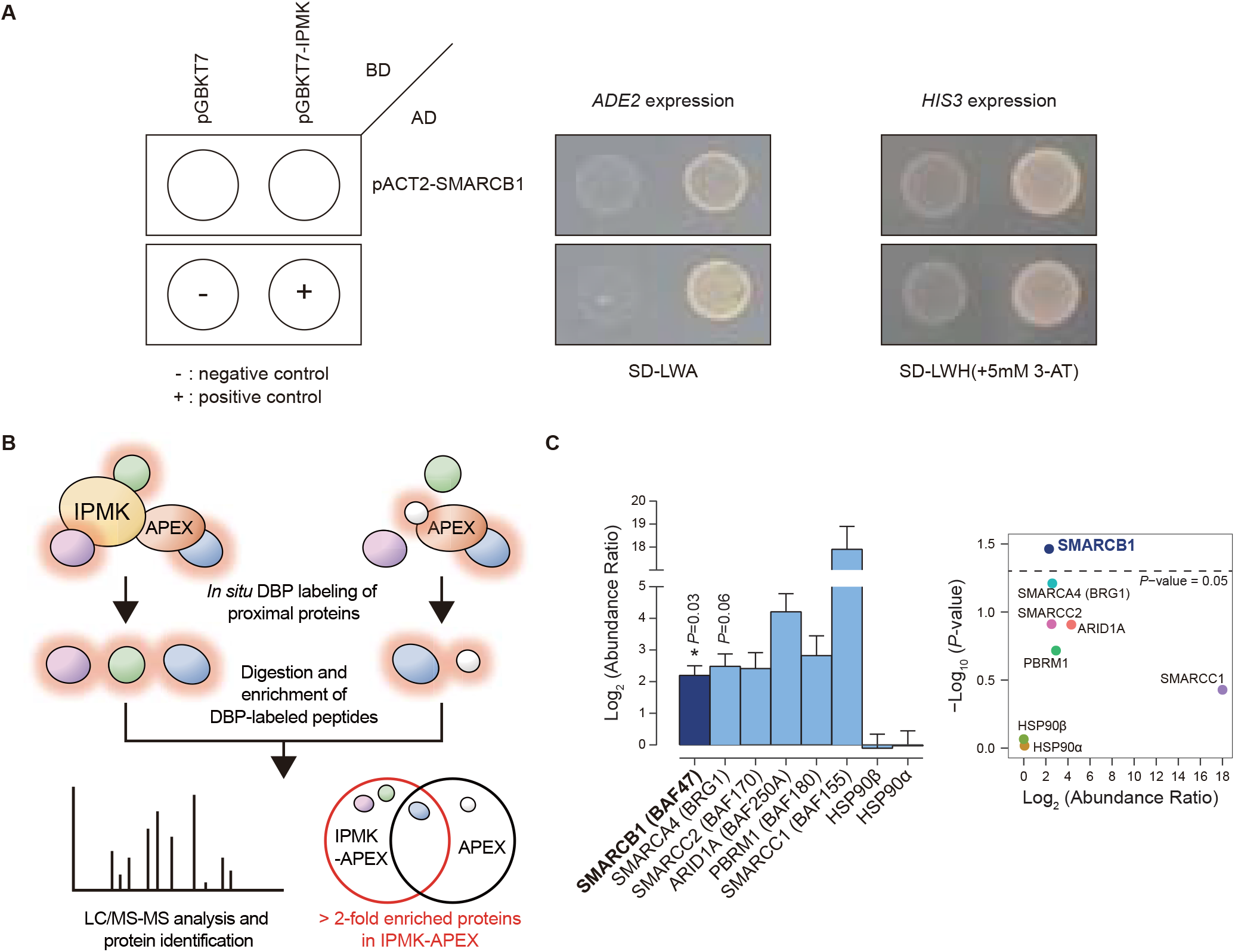
Identification of SMARCB1 as an IPMK-interacting protein via unbiased screening assays. (A) IPMK and SMARCB1 interaction test in yeast strain AH109, containing two reporter genes (*ADE2* and *HIS3*). Yeast cells were co-transformed with either the GAL4-BD fusion plasmid pGBKT7 or pGBKT7-IPMK and the GAL4-AD fusion plasmid pACT2-SMARCB1. The yeast cells were spread on a selection medium lacking leucine and tryptophan (SD-LW) to select co-transformants of bait and prey vectors. Specific interactions between bait and prey proteins were monitored by cell growth on a selection medium lacking leucine, tryptophan, adenine (SD-LWA), or a selection medium lacking leucine, tryptophan, histidine (SD-LWH). 3-AT (3-amino-1,2,4-triazole) was used to suppress leaky *HIS3* expression in transformants to obtain an accurate phenotype. Polypyrimidine tract binding protein (PTB) gene fused with the GAL4 DNA binding domain (BD-PTB) and PTB gene fused with the GAL4 activation domain (AD-PTB) were used as positive controls of bait and prey vectors, respectively. The negative control is the cells transformed with parental bait vector (pGBKT7) and prey vector (pACT2). (B) A schematic diagram displaying identification strategy of IPMK-proximal/interacting proteins, which are biotinylated by APEX-tagged IPMK. (C) Bar graphs showing the relative abundance of biotinylated proteins related to SWI/SNF complex and two negative controls (left). Target proteins were arranged according to their significance (*P*-value, left: significant; right: not significant). A volcano plot showing the relative abundance and significance (*P*-value) of biotinylated proteins related to SWI/SNF complex and two negative controls (right). A dotted line within the volcano plot indicates the *P*-value = 0.05. The relative abundance (abundance ratio) was derived by comparing the fold enrichment of target proteins in IPMK-APEX2-expressed to APEX2-expressed HEK293 cells. *P*-value was calculated using Student’s t-test.

To further identify potential target proteins associating with IPMK in mammalian cells, we performed an *in vivo* proximity-labeling approach using an engineered variant of ascorbate peroxidase (APEX2) fused to IPMK (APEX2-mediated proximity labeling). The IPMK-APEX2 or APEX2 neighboring proteins were biotinylated, enriched with streptavidin beads, and analyzed via mass spectrometry (Figure 1B). A total of 455 IPMK-associated candidate proteins was identified by comparing neighboring proteins of APEX2-IPMK with those of APEX2 only (background, used as a negative control) using two-fold enrichment score differences. Interestingly, by performing ConsensusPathDB (Herwig, Hardt, Lienhard, & Kamburov, 2016) using the IPMK-associated candidates, we detected the enriched protein complex-based sets related to BRG1-, BAF-, or SWI/SNF complex-associated complexes (Supplementary Table 2). Notably, among these candidates, we detected SWI/SNF complex-associated factors, including SMARCB1 (BAF47), BRG1 (SMARCA4), SMARCC2 (BAF170), ARID1A (BAF250A), PBRM1 (BAF180), and SMARCC1 (BAF155), as IPMK-proximal/binding target proteins (Figure 1C). Furthermore, among these SWI/SNF complex subunits, we found that SMARCB1 most significantly interacts with or being proximal to IPMK (Figure 1C), consistent with the yeast two-hybrid screening results (Figure 1A). Intriguingly, we also detected core histones (histone H2B, H3.1, and H4) as IPMK-proximal/binding targets (Figure 1—figure supplement 1), supporting that IPMK binds/proximal with SWI/SNF complex, which binds to nucleosomes *in vivo*. Collectively, these results from two unbiased screening experiments (including yeast two-hybrid screening assay and APEX2-mediated proximity-labeling-based proteomics) strongly indicate that IPMK physically associates with the SWI/SNF complex.

### Physical interaction between IPMK and core subunits of SWI/SNF complex

To confirm the physical interaction between IPMK and SMARCB1, we first performed an *in vitro* binding assay using recombinant IPMK and SMARCB1 proteins. Notably, we detected a direct protein-protein interaction between IPMK and SMARCB1 *in vitro* (Figure 2A, see lane 2 and lane 4). To confirm the physical association between IPMK and core subunits of SWI/SNF (BAF) complex, we performed binary protein interactions assays with baculovirus-mediated expression. We co-infected Sf9 insect cells with baculoviruses expressing FLAG-IPMK and untagged individual subunits of SWI/SNF complex, including SMARCB1, BRG1, BAF155 (SMARCC1), and BAF170 (SMARCC2). Then, we performed FLAG M2 agarose immunoprecipitation and immunoblotting. Importantly, we detected direct protein-protein interactions between IPMK-SMARCB1, IPMK-BRG1, and IPMK-BAF155 *in vitro*, while we did not detect IPMK-BAF170 interactions (Figure 2B, see lane 7 and lane 8). Together, these results indicate that IPMK directly/individually binds to SMARCB1, BRG1, and BAF155 *in vitro*.

**Figure 2.**
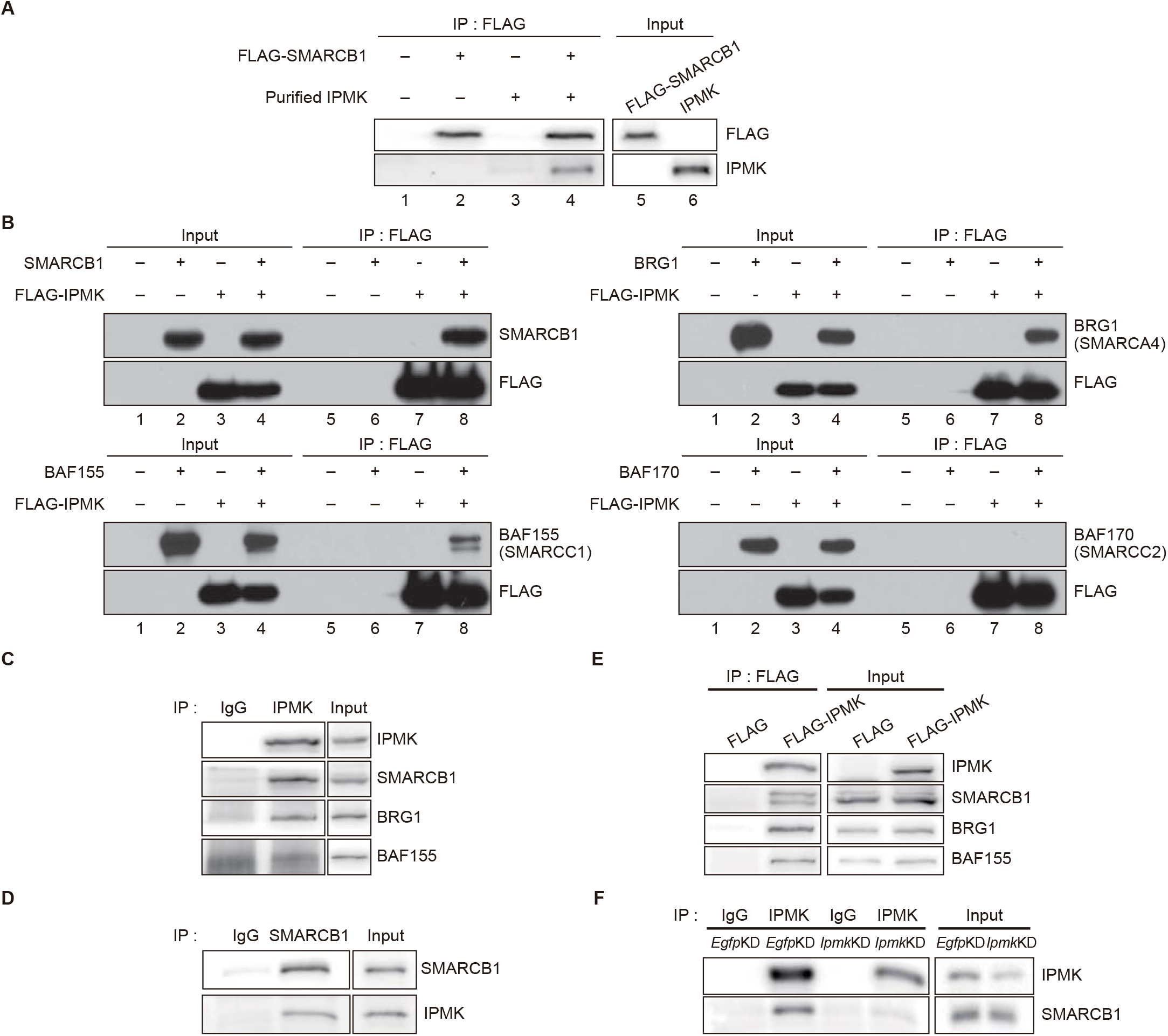
IPMK binds to SMARCB1 and other components of the SWI/SNF complex. (A) Purified IPMK and *in vitro* translated FLAG-SMARCB1 were co-incubated, immunoprecipitated with FLAG antibody, and subjected to immunoblotting. (B) Sf9 insect cells were co-infected with baculoviruses expressing FLAG-IPMK and individual subunits of SWI/SNF complex (SMARCB1, BRG1, BAF155, and BAF170), followed by FLAG M2 agarose immunoprecipitation and immunoblotting. (C) IPMK and IgG were immunoprecipitated from E14Tg2a cells and subjected to immunoblotting. (D) SMARCB1 and IgG were immunoprecipitated from E14Tg2a cells and subjected to immunoblotting. (E) E14Tg2a cells were transfected with FLAG-IPMK or FLAG (a control vector), followed by FLAG immunoprecipitation and immunoblotting. (F) E14Tg2a cells were transfected with siRNA against *Egfp* (*Egfp*KD) and *Ipmk* (*Ipmk*KD), immunoprecipitated with IPMK and IgG, and subjected to immunoblotting.

To investigate whether these IPMK-SMARCB1/BRG1/BAF155 interactions also occurred *in vivo*, we performed co-immunoprecipitation experiments using mammalian cells. Importantly, we detected a direct association of endogenous IPMK and SMARCB1 in mouse embryonic stem cells (mESCs) (Figure 2C and D) and in mouse embryonic fibroblasts (MEFs) (Figure 2—figure supplement 1A–C). Consistent with our *in vitro* binding assays (Figure 2A and B), we detected SMARCB1, BRG1, and BAF155 in endogenous IPMK immunoprecipitates (Figure 2C) and FLAG-IPMK overexpressed immunoprecipitates (Figure 2E) in mESCs. Intriguingly, we detected SMARCB1, BRG1, and BAF170 in endogenous IPMK immunoprecipitates in MEFs (Figure 2—figure supplement 1A and B). To determine the specificity of the IPMK-SMARCB1 physical interaction, RNAi-mediated knockdown of *Ipmk* (*Ipmk*KD) and *Smarcb1* (*Smarcb1*KD) was conducted in mESCs and MEFs. We first confirmed the successful knockdown of both *Ipmk* and *Smarcb1* by quantifying the protein levels (Figure 2F, Figure 2—figure supplement 1D and E). In addition, we observed that *Ipmk* knockdown did not affect the protein levels of SWI/SNF complex subunits, and *Smarcb1* knockdown did not affect the protein levels of IPMK both in mESCs and MEFs (Figure 2—figure supplement 1D). Importantly, in IPMK immunoprecipitates, a significant reduction in SMARCB1 signals was observed in *Ipmk*KD mESCs compared to the control (*Egfp*KD) mESCs (Figure 2F). We also found that SMARCB1 and BRG1 signals in IPMK immunoprecipitates were significantly decreased upon *Smarcb*KD MEFs compared to the control (*Egfp*KD) MEFs (Figure 2—figure supplement 1E). Lastly, in SMARCB1 immunoprecipitates, a significant reduction of IPMK signals was detected in IPMK-null MEFs (Figure 2—figure supplement 1C). Together, these results indicate that IPMK directly associates with the core subunits of SWI/SNF complex *in vivo* (IPMK-SMARCB1/BRG1/BAF155 in mESCs; IPMK-SMARCB1/BRG1/BAF170 in MEFs; IPMK-SMARCB1 binding is specific and observed both in mESCs and MEFs).

Considering our previous results (unbiased screenings, *in vitro*, and *in vivo* immunoprecipitation assays), it is highly plausible that IPMK physically associates with SWI/SNF complex. To confirm this, we first performed a co-immunoprecipitation assay by overexpressing IPMK and SMARCB1 in human embryonic kidney (HEK)-293T cells. Consistently, we observed a physical interaction between IPMK and SMARCB1 (Figure 2—figure supplement 1F). Next, we performed GST (-IPMK) pull-down assays by overexpressing GST-IPMK or GST alone in HEK293T cells. Notably, compared to GST alone, we detected core subunits of SWI/SNF (BAF)/PBAF complexes, including SMARCB1, BAF155, BAF170, PBRM1, BAF250A, and BRM, in GST-IPMK pulled-down samples (Figure 2—figure supplement 1G), as consistent with our APEX2-mediated proximity labeling assays (Figure 1C). Lastly, we purified native SWI/SNF (BAF) complexes from FLAG-DPF2 cell lines (Figure 2—figure supplement 1H), co-incubated the purified native SWI/SNF complexes, and purified GST-IPMK or GST alone, then performed GST pull-down assays. As expected, we detected the core subunits of SWI/SNF complexes (including SMARCB1, BRG1, BAF155, and BAF170) in GST-IPMK pulled-down samples (Figure 2—figure supplement 1I). Collectively, our combined results (unbiased screenings, *in vitro*/*in vivo* immunoprecipitation, and *in vitro* pull-down assays) strongly imply that IPMK physically associates with mammalian SWI/SNF complexes by directly binds to SMARCB1, BRG1, and BAF155 (SMARCC1).

### Mapping the reciprocal binding sites of IPMK and SMARCB1

Among three IPMK-binding proteins (SMARCB1, BRG1, and BAF155), SMARCB1 exhibited the most robust interaction with IPMK (Figure 1 and 2). Regarding this, we conducted yeast two-hybrid assays to identify the specific binding domains of SMARCB1 required for interacting with IPMK. Various prey vectors encoding different SMARCB1 domains were cloned, and two-hybrid analyses were performed. Interestingly, the prey vectors expressing amino acids of 99-245, 99-319, and full-length SMARCB1 showed positive signals in the two-hybrid system (Figure 3—figure supplement 1A), indicating that the Rpt1 and Rpt2 domains of SMARCB1 participate in the protein-protein interactions of SMARCB1 and IPMK.

To further dissect the reciprocal binding sites required for SMARCB1-IPMK physical binding, various SMARCB1 constructs were designed and overexpressed in HEK293T cells. Based on the domain map of SMARCB1 (Figure 3A, top), we first designed SMARCB1 constructs with C-terminal deletion (Figure 3A, middle). We confirmed that the Rpt1 domain of SMARCB1 is essential for IPMK interaction by immunoprecipitating the overexpressed deletion constructs (Figure 3B, compare lane 3 to lane 2, 4, and 5). We then generated additional N-terminal deleted constructs of SMARCB1 (Figure 3A, bottom). Consistent with the results of yeast two-hybrid assays (Figure 3—figure supplement 1A), we observed that the constructs containing the Rpt1 or Rpt2 domains associate with IPMK (Figure 3C, lanes 2, 3, 4, and 5 show positive immunoblotting signals). By independently overexpressing each SMARCB1 domain, we found that both the Rpt1 and Rpt2 domains of SMARCB1 could bind to IPMK (Figure 3D and E, see lane 3, 4, and 6). Additionally, we dissected Rpt1 and Rpt2 domains into β sheets and α helices based on their structures. In Rpt1, two β sheets and two helices are required for IPMK-binding (Figure 3—figure supplement 1B, compare lane 4 to lane 2 and 3). In Rpt2, only two β sheets but not α helices were bound to IPMK (Figure 3—figure supplement 1C, compare lane 2 and lane 3). These observations (for IPMK-binding, both sheets and helices of Rpt1 or/and only sheets of Rpt2 are required) were supported by overexpressing combinations of Rpt1 and Rpt2 domains (Figure 3—figure supplement 1D, lanes 3 and 4 show positive signals). Lastly, this was further confirmed by the fact that SMARCB1 lacking Rpt1 and Rpt2 could not bind to IPMK (Figure 3—figure supplement 1E, compare lane 2 and lane3, and S3F). Taken together, we concluded that the sheets/helices of Rpt1 and sheets of Rpt2 are the major IPMK interacting sites in SMARCB1.

**Figure 3.**
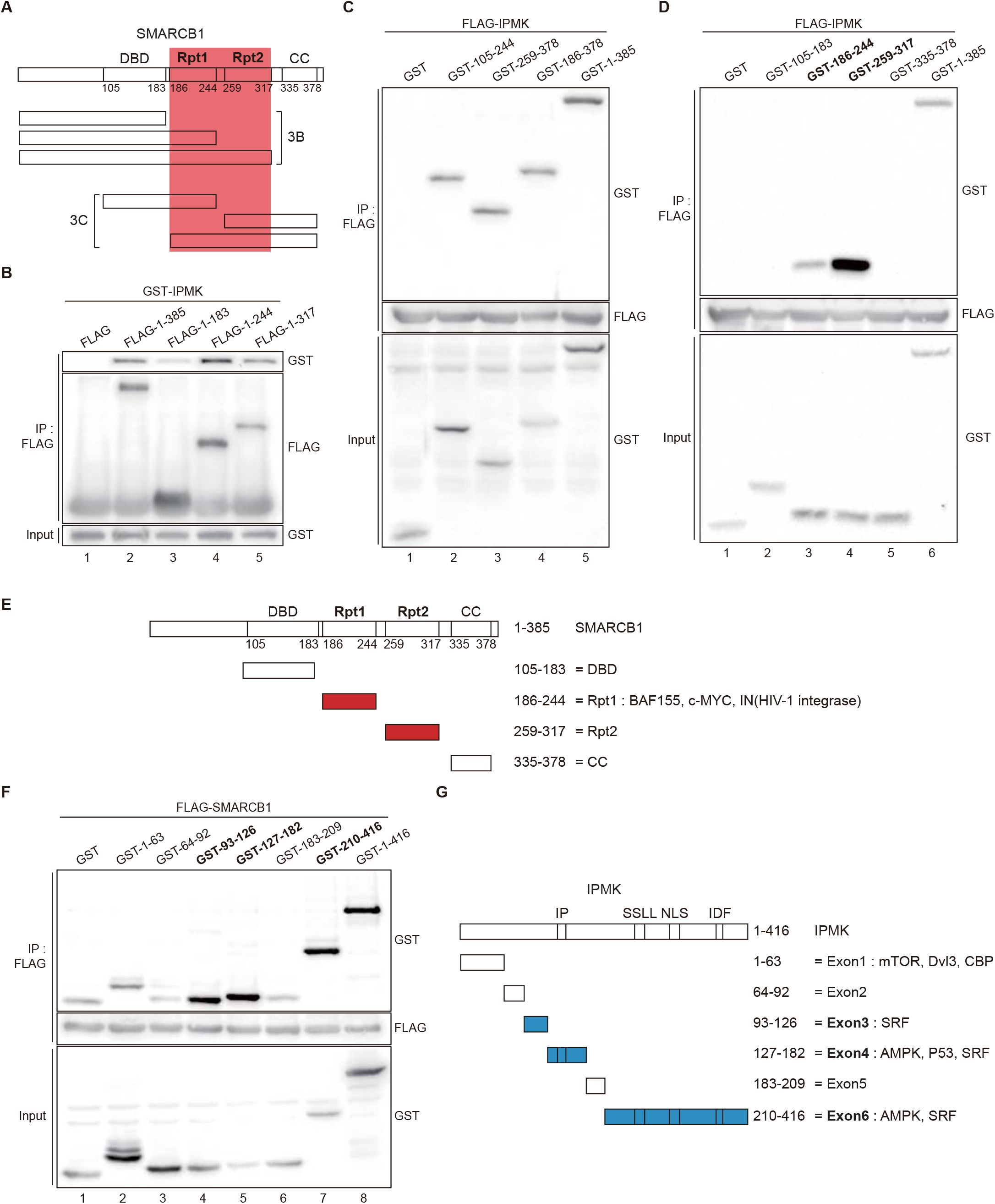
Identification of distinct domains required for IPMK-SMARCB1 interactions. (A) A schematic diagram of the human SMARCB1 fragments used for the binding studies (B and C). The IPMK-binding sites (Rpt1 and Rpt2) are highlighted in red. (B) HEK293T cells were co-transfected with GST-IPMK and FLAG (a control vector) or FLAG-SMARCB1 fragments, followed by immunoprecipitation with FLAG antibody, and subjected to immunoblotting. (C and D) HEK293T cells were co-transfected with FLAG-IPMK and GST (a control vector) or GST-SMARCB1 fragments, followed by immunoprecipitation with FLAG antibody, and subjected to immunoblotting. The specific IPMK-binding SMARCB1 fragments are highlighted in bold. (E) A schematic diagram of the human SMARCB1 domains. SMARCB1 fragments used for the binding studies (D) are indicated below with the numbers of amino acid sequences. The specific IPMK-binding SMARCB1 fragments (Rpt1 and Rpt2) are highlighted in red. (F) HEK293T cells were co-transfected with FLAG-SMARCB1 and GST (a control vector) or GST-IPMK fragments, followed by immunoprecipitation with FLAG antibody, and subjected to immunoblotting. (G) A schematic diagram of human IPMK domains. IPMK fragments used for the binding studies (F) are indicated below with the numbers of amino acid sequences. Key domains for inositol binding (IP), kinase activity (SSLL and IDF), and nuclear localization signal (NLS) are depicted. The specific SMARCB1-binding IPMK fragments (Exon3, 4, and 6) are highlighted in blue.

Reciprocally, to identify which IPMK domains are required for SMARCB1 binding, we designed several GST-tagged constructs of IPMK (Figure 3G) and conducted immunoprecipitation experiments. Intriguingly, we found that IPMK-SMARCB1 binding was primarily mediated by three IPMK regions, including exon 3, exon 4, and exon 6 (Figure 3F, lanes 4, 5, 7, and 8 show positive signals, and 3G), which comprises the inositol binding site and the kinase domain. Taken together, our results elucidated the specific reciprocal biding sites of IPMK-SMARCB1 interaction.

### Co-localization of IPMK and BRG1

Our results demonstrated that IPMK directly binds to the core subunits of mammalian SWI/SNF complexes (SMARCB1, BRG1, and BAF155) and physically associates with native SWI/SNF complexes. Regarding this, one can speculate that IPMK may play an important role in chromatin regulation. However, the region where this IPMK-SWI/SNF interaction occurs *in vivo* and the detailed localization and the role of IPMK in the chromatin remains elusive. To decipher these issues, we first conducted a chromatin fractionation assay using mESCs and MEFs. Notably, we found that IPMK is evenly distributed in all three fractions (cytoplasm, nucleoplasm, and chromatin), whereas SMARCB1 and BRG1 primarily reside in the chromatin fraction both in mESCs and MEFs (Figure 4A and Figure 4—figure supplement 1A-C). To further investigate whether IPMK and SMARCB1 expressions affect each other’s distribution, RNAi-mediated knockdown of *Ipmk* (*Ipmk*KD) and *Smarcb1* (*Smarcb1*KD) was conducted before the chromatin fractionation assay. We observed that the distribution of SMARCB1 was unaffected by *Ipmk*KD (Figure 4A), and the distribution of IPMK was unaffected by *Smarcb1*KD (Figure 4—figure supplement 1C). Together, these results indicate that IPMK, BRG1, and SMARCB1 reside together in the chromatin.

**Figure 4.**
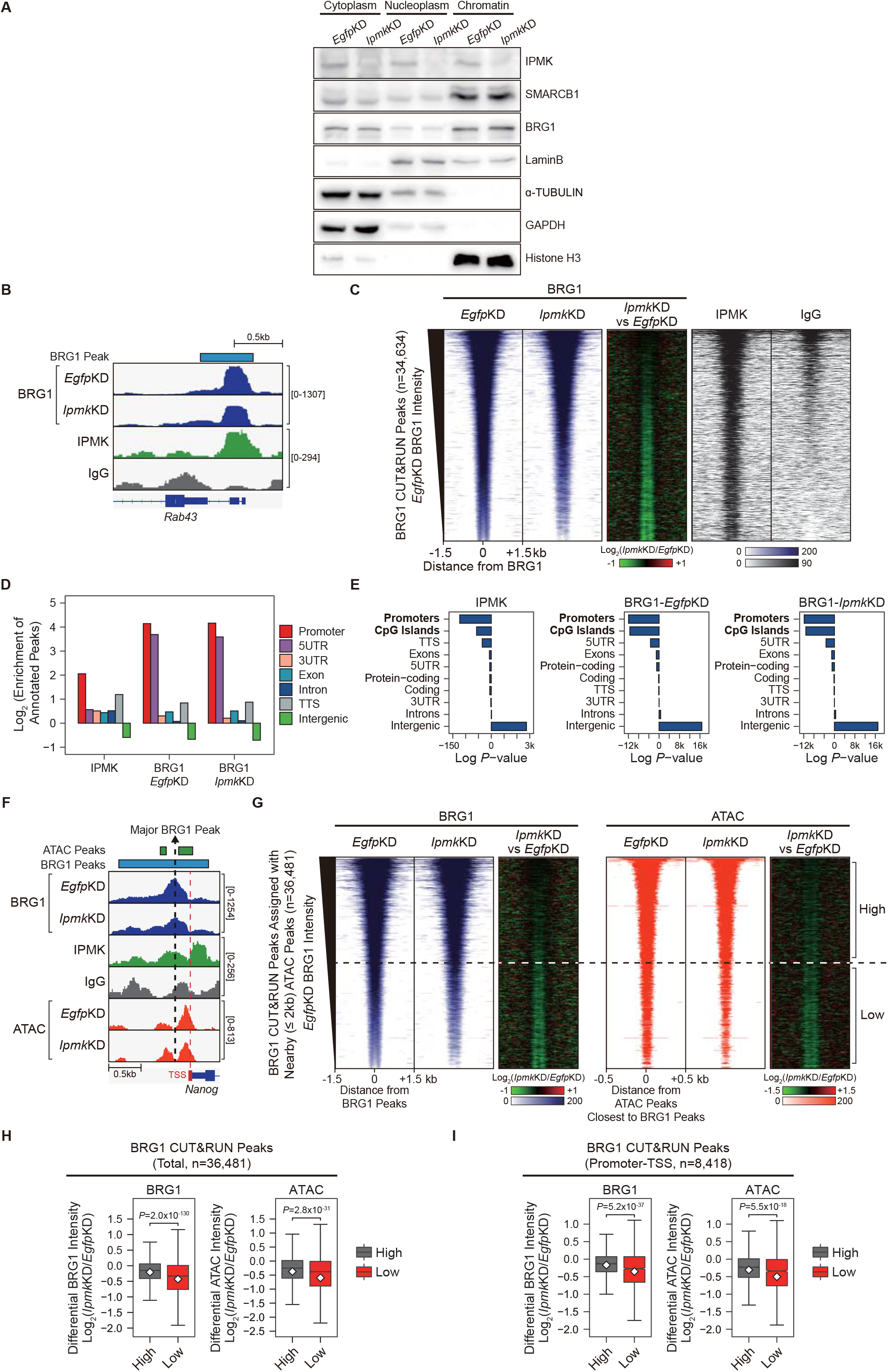
IPMK-BRG1 are co-localized at promoter-TSS, and IPMK regulates BRG1 localization. (A) E14Tg2a cells were transfected with siRNA against *Egfp* (*Egfp*KD) and *Ipmk* (*Ipmk*KD) and then fractionated into the cytoplasm, nucleoplasm, and chromatin fractions. Immunoblotting with IPMK, SMARCB1, BRG1, and fractionation markers was then performed. (B) Examples of CUT&RUN assays in E14Tg2a cells, including representative results for BRG1 (*Egfp*KD and *Ipmk*KD), IPMK, and IgG. The BRG1 CUT&RUN peak (*Egfp*KD cells) is marked as a blue box on top. (C) Heatmaps representing CUT&RUN results for BRG1 (*Egfp*KD, *Ipmk*KD, and their comparison), IPMK, and IgG at BRG1 CUT&RUN peaks (*Egfp*KD cells) as indicated on top. All heatmaps were aligned at 34,634 BRG1 CUT&RUN peaks (rows) and sorted in descending order by the BRG1 intensity of *Egfp*KD cells. (D) Bar graphs showing the Log_2_ enrichment of CUT&RUN peaks (IPMK and BRG1-*Egfp*KD, -*Ipmk*KD) annotated with various regions of the mouse genome. (E) Bar graphs showing the significance (Log *P*-value) of CUT&RUN peaks (IPMK and BRG1-*Egfp*KD, -*Ipmk*KD) annotated with various regions of the mouse genome. For each CUT&RUN peak, genome annotations (e.g., promoters or CpG islands) are sorted in descending order according to their significance (*P*-values, top-bottom, whereas top-more significant and bottom-less significant). (F) Examples of BRG1 (*Egfp*KD and *Ipmk*KD), IPMK, IgG CUT&RUN, and ATAC-seq (*Egfp*KD and *Ipmk*KD) assays in E14Tg2a cells. The BRG1 CUT&RUN peak (*Egfp*KD cells) and ATAC-seq peaks are marked as blue and green boxes on top. Major BRG1 peak (the most enriched site) and transcription start site (TSS) are indicated as black and red dotted lines. (G) Heatmaps representing BRG1 CUT&RUN (*Egfp*KD, *Ipmk*KD, and their comparison) at BRG1 CUT&RUN peaks (*Egfp*KD cells) assigned with nearby (within 2kb) ATAC-seq peaks (left). BRG1 peaks without nearby ATAC-seq peaks were excluded. In order to match the arrangement with ATAC-seq peaks (right), a BRG1 peak containing multiple ATAC-seq peaks was included without deduplication. Heatmaps representing ATAC-seq signals (*Egfp*KD, *Ipmk*KD, and their comparison) at ATAC-seq peaks assigned with closest BRG1 CUT&RUN peaks that were used for heatmaps on the left (right). All heatmaps were aligned at 36,481 BRG1 CUT&RUN peaks (left) or 36,481 ATAC-seq peaks (right) and sorted in descending order by the BRG1 intensity of *Egfp*KD cells. High and Low groups were divided equally (n=18240 and 18241, respectively) according to the BRG1 intensity of *Egfp*KD cells. (H) Box plots showing the differential BRG1 (left) and ATAC (right) intensity upon *Ipmk*KD at High (grey) and Low (red) BRG1 CUT&RUN peaks (left) and corresponding (closest) ATAC-seq peaks (right). High and Low groups (n=18240 and 18241, respectively) were divided according to the BRG1 intensity of *Egfp*KD cells. (I) Box plots showing the differential BRG1 (left) and ATAC (right) intensity upon *Ipmk*KD at High (grey) and Low (red) BRG1 CUT&RUN peaks localized at Promoter-TSS (left) and corresponding (closest) ATAC-seq peaks (right). High and Low groups (n=5,800 and 2,618, respectively) were derived from (G and H). (H and I) *P*-values were calculated using the Wilcoxon rank sum test.

We then sought to determine where this event (IPMK, BRG1, and SMARCB1 localizing at the chromatin) takes place within the chromatin. To investigate the localization of BRG1 and IPMK within the chromatin, we performed CUT&RUN (cleavage under targets and release using nuclease) assays (Skene & Henikoff, 2017) in mESCs. In accordance with our results of the IPMK-SWI/SNF complex’s physical association, we found that IPMK was co-localized with BRG1 in a genome-wide manner (Figure 4B and C). Next, we performed peak annotation to analyze the genomic regions (e.g., promoters or intergenic regions) enriched with CUT&RUN peaks. Given that BRG1, a catalytic subunit of the SWI/SNF complex, is known to localize at the promoter-transcription start site (TSS) (de Dieuleveult et al., 2016), we confirmed that BRG1 was significantly enriched at promoters of the mouse genome (Figure 4D and E). Notably, we detected that IPMK was also significantly enriched at promoters (Figure 4D and E). Collectively, these results strongly indicate that IPMK and BRG1 are co-localized at the chromatin, particularly at the promoter region, which further support our previous results of physical association between IPMK and SWI/SNF complex.

### IPMK regulates the BRG1 occupancy and impacts the BRG1-mediated chromatin accessibility

To further elucidate the role of IPMK in BRG1 localization, we performed BRG1 CUT&RUN assays upon *Ipmk*KD and compared them to *Egfp*KD (control) in mESCs. Interestingly, we observed a decreased BRG1 occupancy upon *Ipmk*KD at specific promoter-TSS regions with enriched IPMK (Figure 4B and F). Strikingly, we found that the genome-wide BRG1 localization was severely disrupted (decreased BRG1 occupancy) upon *Ipmk*KD at BRG1 CUT&RUN peaks with low BRG1 enrichment in *Egfp*KD mESCs (bottom half of the heatmaps, termed as Low) (Figure 4C, G, and H). In addition, we detected that BRG1 CUT&RUN peaks’ genomic distributions were unaffected by *Ipmk*KD (Figure 4D and E), suggesting that *Ipmk*KD do not affect the global distribution (changes in peak positions) of BRG1 but impacts the global occupancy of BRG1.

It is previously known that BRG1 regulates chromatin accessibility at NFR (nucleosome free regions) of TSS in mESCs (de Dieuleveult et al., 2016). To investigate the effect of *Ipmk*KD-induced decreased BRG1 occupancy on chromatin accessibility, we performed ATAC-seq (assay for transposase-accessible chromatin using sequencing) upon *Ipmk*KD and compared them to *Egfp*KD (control) in mESCs. Notably, we observed that both ATAC-seq signals and BRG1 occupancy were reduced upon *Ipmk*KD at promoter-TSS regions of *Nanog* (Figure 4F). Furthermore, we observed some discrepancies in major peak positions when comparing BRG1 CUT&RUN peaks (*Egfp*KD, termed as BRG1 peaks) and ATAC-seq peaks (*Egfp*KD, termed as ATAC peaks) (Figure 4F). To precisely assess the effect of *Ipmk*KD-induced decreased BRG1 occupancy on chromatin accessibility, we assigned ATAC peaks to the nearby (within 2kb) BRG1 peaks and selected these BRG1 peaks for further analysis (we excluded BRG1 peaks without nearby ATAC peaks) (Figure G). In addition, BRG1 peaks containing or assigned with multiple ATAC peaks were included without deduplication to match the same ordering as ATAC peaks (the same ordering – alignment of heatmaps’ row – was applied for BRG1 and ATAC peaks in Figure 4G). As expected, we observed that the global BRG1 occupancy was reduced upon *Ipmk*KD at BRG1 peaks with low BRG1 intensity (bottom half of the heatmaps, termed as Low) (Figure G and H), consistent with our previous observations (Figure 4C). Surprisingly, at Low BRG1 peaks (Low), we observed that both BRG1 occupancy and BRG1-mediated chromatin accessibility (ATAC-seq signals closest to the BRG1 CUT&RUN peaks) were significantly reduced upon *Ipmk*KD in a genome-wide manner (Figure 4G and H). Furthermore, the reduced BRG1 and BRG1-associated chromatin accessibility (ATAC-seq) upon *Ipmk*KD were also detected at promoter-TSS regions (Figure 4I), consistent with the previous study (BRG1 primarily maintains chromatin accessibility at promoter-TSS regions) (de Dieuleveult et al., 2016). Taken together, these results indicate that IPMK regulates the global BRG1 occupancy and corresponding BRG1-mediated chromatin accessibility in mESCs.

### IPMK plays an important role in BRG1 localization and chromatin accessibility at promoter-TSS

We next focused on the promoter-TSS regions, where BRG1 and IPMK were significantly enriched (Figure 4D and E). To further dissect the genome-widely decreased BRG1 occupancy upon *Ipmk*KD, we classified the distinct clusters of promoter-TSS based on the promoter types and changes in BRG1 intensity at defined regions relative to TSS positions (Figure 5C-G). Initially, by analyzing MNase-seq (chromatin digestion with micrococcal nuclease combined with sequencing) and ATAC-seq data in mESCs (*Egfp*KD), we observed nucleosome-depleted/chromatin accessible regions, known as NFR (nucleosome free regions), and detected −1 and +1 nucleosomes near TSS (Figure 5A, left). We also confirmed that BRG1 was abundant at TSS by analyzing BRG1 CUT&RUN assays (Figure 5A, left). Based on the signal intensity of ChIP-seq (chromatin immunoprecipitation sequencing) against histone H3K4me3 and H3K27me3, we classified promoters into three types: H3K4me3-Low, H3K4me3-Only (high H3K4me3, low H3K27me3), and Bivalent (high H3K4me3, high H3K27me3). Considering nucleosome intensity and ATAC-seq signals, we detected that chromatin was highly accessible in H3K4me3-Only, moderately accessible in bivalent, and inaccessible in H3K4me3-Low promoters (Figure 5A, right). Next, we categorized the 9,042 TSS with decreased BRG1 occupancy upon *Ipmk*KD (1.5 fold changes in BRG1 occupancy compared to *Egfp*KD, see also Figure 5D, left Total) into the three promoter types (Figure 5B). Since our goal was to examine the *Ipmk*KD-induced ‘decreased’ BRG1 occupancy (changes from enriched to depleted BRG1 signal upon *Ipmk*KD as seen in Figure 4C and G-I), we excluded the H3K4me3-Low promoters, which exhibited the extremely low BRG1 signals (Figure 5A, top right). Furthermore, regarding the fact that TSS divided the BRG1 intensity into two (Figure 5A, top left), we defined two genomic regions (Upstream and Downstream) that coincided with the two major BRG1 intensity (which also coincided with −1 and +1 nucleosomes), respectively (Figure 5A, top). We then categorized the 9,042 TSS with *Ipmk*KD-induced decreased BRG1 occupancy into five clusters based on the combinatorial changes in BRG1 levels upon *Ipmk*KD at the previously defined two genomic regions (Figure 5C, left and middle) and subdivided these five clusters based on two promoter types (Figure 5C, right). Regarding the number of promoter-TSS with decreased BRG1 level upon *Ipmk*KD, H3K4me3-Only promoters (Figure 5B and C, right) and Cluster2/3 (Figure 5C) occupied a large proportion. In accordance with our defined classification (Figure 5C, left), we detected that BRG1 occupancy was reduced upon *Ipmk*KD at both promoters and five distinctive clusters (Figure 5D).

**Figure 5.**
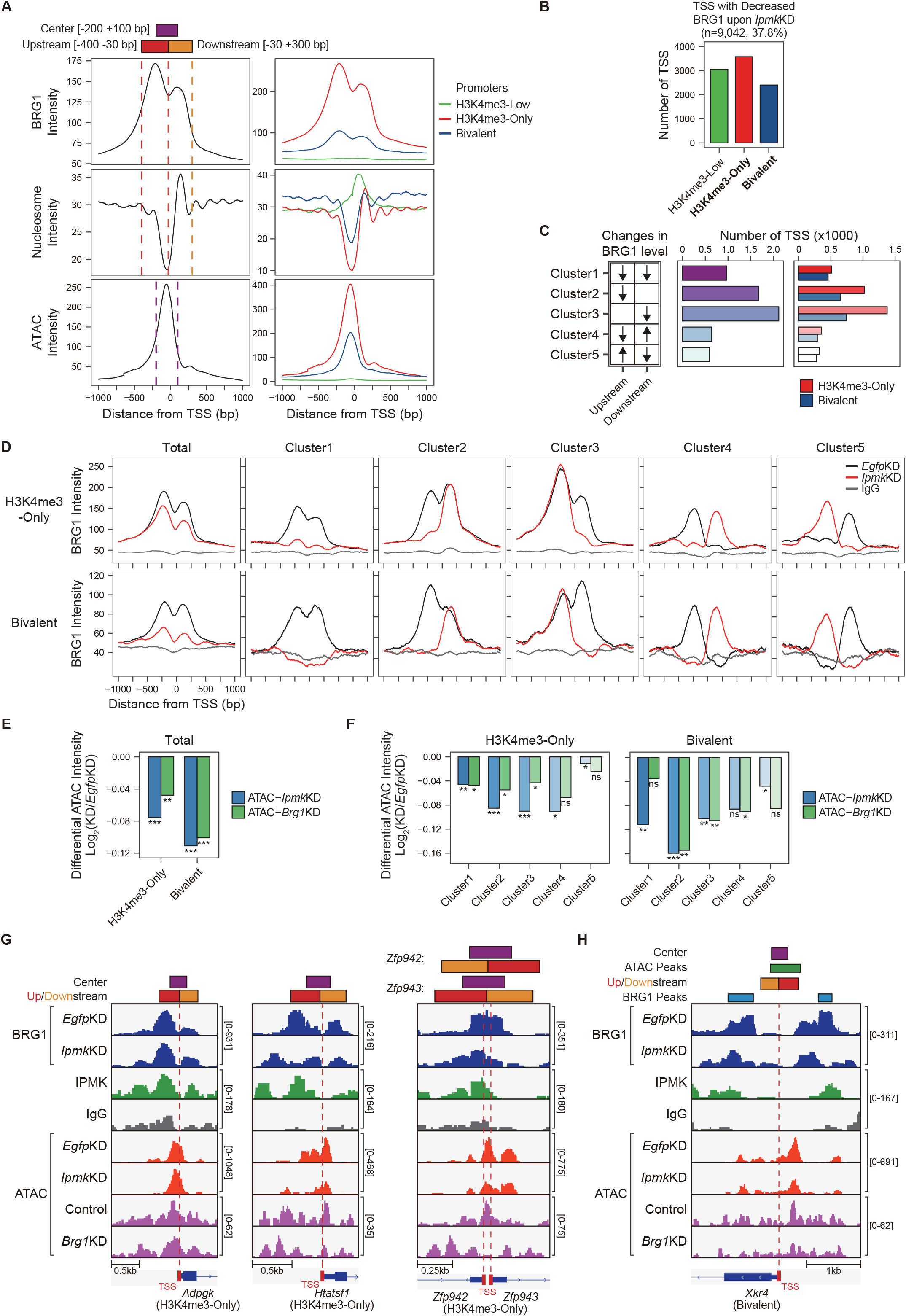
IPMK affects chromatin accessibility at promoter-TSS by regulating the BRG1 localization. (A) Line plots showing the average enrichments of BRG1, nucleosome (MNase-seq, GSM5253962 and GSM5253963), and ATAC-seq signals (ATAC) at TSS of total genes (left) and TSS of three promoter types (right). Three genomic regions, indicated on top (see also dotted lines on left), were defined according to the relative position of enriched ATAC-seq intensity (Center, purple) and enriched BRG1 intensity (Upstream and Downstream, red and orange, respectively). Green, red, and blue lines on the right indicate H3K4me3-Low, H3K4me3-Only, and bivalent promoters, respectively. (B) Bar graphs showing the number of TSS (TSS exhibiting decreased BRG1 intensity upon *Ipmk*KD) with different promoter types. (C) A diagram displaying five clusters of TSS classified by changes in BRG1 level at Up/Downstream regions (defined in (A)) upon *Ipmk*KD (left). Arrows with downwards and upwards indicate decreased and increased BRG1 level upon *Ipmk*KD, respectively. Bar graphs showing the number of five TSS clusters (middle) with different promoter types (right). (D) Line plots showing the average enrichments of BRG1 (*Egfp*KD and *Ipmk*KD) and IgG at TSS with two promoter types (see the total on left, H3K4me3-Only and bivalent promoters are shown on top and bottom, respectively) and with five TSS clusters. Black, red, and gray lines indicate BRG1 intensity upon *Egfp*KD, *Ipmk*KD, and IgG intensity, respectively. (E) Bar graphs showing the average of differential ATAC-seq intensity (Log_2_ KD/*Egfp*KD) upon *Ipmk*KD (blue) and *Brg1*KD (green) at TSS with two promoter types. (F) Bar graphs showing the average of differential ATAC-seq intensity (Log_2_ KD/*Egfp*KD) upon *Ipmk*KD (blue) and *Brg1*KD (green) at five TSS clusters with H3K4me3-Only (left) and bivalent promoters (right). (E and F) *P*-values were derived using Wilcoxon signed rank test (*P < 0.01; **P < 1×10^-4^; ***P < 1×10^-10^; ns, not significant). (G) Examples of BRG1 (*Egfp*KD and *Ipmk*KD), IPMK, IgG CUT&RUN, and ATAC-seq (*Egfp*KD, *Ipmk*KD, Control, and *Brg1*KD) assays at TSS of *Adpgk* (left), *Htatsf1* (middle), and *Zfp942/943* (right). Center and Up/Downstream regions are marked as purple and red/orange boxes on top, respectively. TSS are marked with red boxes (bottom) and dotted lines. (H) Examples of BRG1 (*Egfp*KD and *Ipmk*KD), IPMK, IgG CUT&RUN, and ATAC-seq (*Egfp*KD, *Ipmk*KD, Control, and *Brg1*KD) assays at TSS of *Xkr4*. The BRG1 CUT&RUN peaks (*Egfp*KD cells) and ATAC-seq peaks are marked as blue and green boxes on top. Center and Up/Downstream regions are marked as purple and red/orange boxes on top, respectively. TSS is marked with a red box (bottom) and a dotted line.

Previously, it is reported that BRG1, localized at −1 nucleosome in wide NFR (median length 808 bp) of H3K4me3-Only and bivalent promoters, positively regulates the chromatin accessibility of NFR, whereas BRG1 that localized at +1 nucleosome in narrow NFR (median length 28 bp) of H3K4me3-Only promoters tends to inhibit the chromatin accessibility of NFR in mESCs (de Dieuleveult et al., 2016). To elucidate the effect of *Ipmk*KD-induced decreased BRG1 occupancy on chromatin accessibility at two promoters and five clusters, we analyzed two ATAC-seq data of ours (*Ipmk*KD) and publically released (*Brg1*KD, GSE64825). We calculated the differential ATAC-seq signals (KD vs. controls) at Center regions, where ATAC-seq intensity is highly enriched (Figure 5A). Consistent with our previous genome-wide results (Figure 4G-I), we found that ATAC-seq signals (chromatin accessibility) were significantly reduced upon *Ipmk*KD at two promoters (Figure 5E) and most clusters (Figure 5F), where these promoters/clusters were defined by decreased BRG1 occupancy upon *Ipmk*KD. Notably, we observed that ATAC-seq signals (chromatin accessibility) were decreased similarly upon *Ipmk*KD and *Brg1*KD at two promoters (Figure 5E) and five clusters (Figure 5F). Since the reduced BRG1 occupancy upon *Ipmk*KD partially mimics the *Brg1*KD, the similar result upon *Ipmk*KD and *Brg1*KD further supports that IPMK plays a vital role in chromatin accessibility at promoter-TSS by regulating the BRG1 occupancy. Intriguingly, we found that the ATAC-seq signals at bivalent promoters were more reduced upon *Ipmk*KD compared to those at H3K4me3-Only promoters (Figure 5E and F). Furthermore, we detected that the ATAC-seq signals at Cluster2 of bivalent promoters were more reduced upon *Ipmk*KD than other clusters (Figure 5 F), consistent with the fact that in bivalent promoters, BRG1 is localized at the −1 nucleosome and maintains the chromatin accessibility (de Dieuleveult et al., 2016). Although H3K4me3-Only promoters also contain Cluster2, we did not detect the robust decrease in ATAC-seq signals at Cluster2 of H3K4me3-Only promoters, unlike the bivalent promoter case (Figure 5F). This discrepancy may be due to the different BRG1 occupancy in Upstream (−1 nucleosome) and Downstream (+1 nucleosome) regions of Cluster2 in *Egfp*KD mESCs; BRG1 is highly enriched at Upstream compare to Downstream regions in bivalent promoters, while in H3K4me3-Only promoters, BRG1 levels are relatively similar at both Up/Downstream regions (Figure 5D). Interestingly, although we applied the same criteria when categorizing the five clusters, H3K4me3-Only and bivalent promoters exhibited different BRG1 localizations in *Egfp*KD mESCs at Cluster2 and Cluster3, indicating that these two promoters each possess distinct BRG1 localizations in mESCs. Together, these results suggest that IPMK plays a pivotal role in maintaining the chromatin accessibility of bivalent promoters, particularly by safeguarding the BRG1 occupancy at the −1 nucleosome.

By comparing with the BRG1 unchanged locus (Figure 5G, left), we confirmed the close association between reduced BRG1 level upon *Ipmk*KD and decreased ATAC-seq signals upon *Ipmk*KD and *Brg1*KD at specific loci of H3K4me3-Only and bivalent promoters (Figure 5G and H). However, at a specific locus, we noticed that some BRG1 peaks do not coincide with Up/Downstream regions (Figure 5H), indicating that our promoter-TSS classification using Up/Downstream regions with fixed length does not include promoter-TSS with far away BRG1 peaks. To overcome this, we first selected BRG1 CUT&RUN peaks (*Egfp*KD) that reside in close proximity to TSS (±1 kb) and then newly classified the promoter-TSS into six clusters depending on the position of BRG1 peaks relative to the position of TSS (Figure 5—figure supplement 1A and B). Among six clusters, Cluster1 and 5, which contain BRG1 peaks at TSS and upstream of TSS, respectively, occupied a large proportion (Figure 5—figure supplement 1B). Interestingly, we observed that BRG1 localizations were remarkably similar at Cluster1/2/3, and independently at Cluster4/5 (Figure 5—figure supplement 1A). To simplify the clustering, we merged Cluster1/2/3 as ClusterC and Cluster4/5 as ClusterL (Cluster6 is equivalent to ClusterR) (Figure 5—figure supplement 1C and D). Consistently, ClusterC and L, which contain BRG1 peaks at their TSS and upstream of their TSS, respectively, were predominant (Figure 5—figure supplement 1D). We then divided each cluster depending on their decreased or increased BRG1 level by calculating the differential BRG1 level (*Ipmk*KD vs. *Egfp*KD) at the BRG1 peaks corresponding to TSS (Figure 5—figure supplement 1E), instead of at the defined Up/Downstream regions (Figure 5B and C). Consistent with the genome-wide decrease in BRG1 occupancy at the promoter-TSS (Figure 4I), we found that 60-70% of clusters exhibited decreased BRG1 level upon *Ipmk*KD (Figure 5—figure supplement 1E). Furthermore, most of these *Ipmk*KD-induced decreased BRG1 clusters were identified as H3K4me3-Only promoters (Figure 5—figure supplement 1F).

To elucidate the effect of *Ipmk*KD-induced decreased BRG1 occupancy on chromatin accessibility, we first confirmed a significant reduction in BRG1 occupancy upon *Ipmk*KD at the pre-defined BRG1-decreased clusters/promoters (Figure 5—figure supplement 1G). Next, we calculated differential ATAC-seq signals (*Ipmk*KD vs. *Egfp*KD and *Brg1*KD vs. control) at ATAC-seq peaks (*Egfp*KD) that reside near the TSS (±0.5 kb) of each BRG1-decreased cluster/promoter. Consistent with our previous results (Figure 5E and F), we observed that both *Ipmk*KD and *Brg1*KD induced similarly decreased ATAC-seq levels at both promoters and all three clusters (Figure 5—figure supplement 1H). When comparing H3K4me3-Only and bivalent promoters, we observed that ClusterC exhibited a similar reduction, while ClusterL/R exhibited more reduction at bivalent promoters upon *Ipmk*KD (Figure 5—figure supplement 1H, blue). Similarly, we observed that all three clusters exhibited more reduction at bivalent promoters compared to H3K4me3-Only promoters upon *Brg1*KD (Figure 5—figure supplement 1H, green). Lastly, among bivalent promoters, we detected that ClusterL (very similar to Cluster2 in Figure 5) exhibited the most significant reduction in ATAC-seq signals upon *Ipmk*KD (Figure 5—figure supplement 1H).

Collectively, our two alternative analyses suggest that IPMK depletion cause the most severe impacts on the chromatin accessibility at bivalent promoters, which are strongly associated with the *Ipmk*KD-induced decreased BRG1 occupancy at −1 nucleosomes. Taken together, these findings indicate that IPMK regulates BRG1 occupancy and BRG1-mediated chromatin accessibility at promoter-TSS regions.

### Loss of IPMK partially affects transcription via disrupted BRG1 localization and chromatin accessibility at promoter-TSS

To investigate the effect of *Ipmk*KD-driven disrupted BRG1 occupancy and chromatin accessibility on the transcription, we performed high-throughput mRNA sequencing (mRNA-seq) using mESCs with RNAi-mediated knockdown of *Ipmk* (*Ipmk*KD). We next calculated the differential mRNA expression levels (*Ipmk*KD vs. *Egfp*KD) of genes having the promoters with decreased BRG1/ATAC-seq intensity upon *Ipmk*KD (Figure 5D and E, Total). The mRNA expression of genes with both promoter types was down-regulated upon *Ipmk*KD (Figure 5—figure supplement 1I). Intriguingly, we detected that mRNA expression of genes with bivalent promoters was significantly down-regulated upon *Ipmk*KD, compared to H3K4me3-Only promoters (Figure 5—figure supplement 1I). This discrepancy may arise from our previous results that *Ipmk*KD impacted the bivalent promoter-TSS’s chromatin accessibility more than those of H3K4me3-Only promoters (Figure 5E, F, and Figure 5—figure supplement 1H). To check the decreased BRG1/ATAC-seq-associated down-regulation of gene expressions upon *Ipmk*KD, we performed a real-time quantitative polymerase chain reaction (RT-qPCR), a conventional method to check the gene expression. We first determined the successful knockdown of *Ipmk* (Figure 6A). Next, we confirmed that mRNA expression of genes, exhibiting decreased BRG1 occupancy upon *Ipmk*KD and reduced ATAC-seq signals upon *Ipmk*KD/*Brg1*KD at H3K4me3-Only (*Nmral1*) or bivalent (*Phactr3*) promoters, were significantly down-regulated upon *Ipmk*KD (Figure 6B). To examine this in a genome-wide manner, we identified differentially expressed genes (DEGs) by comparing the gene expression of *Ipmk*KD with that of control (*Egfp*KD) cells and identified 300 down-regulated DEGs (Figure 6C). Notably, we observed that BRG1 occupancy and BRG1-mediated chromatin accessibility (ATAC-seq signals) were both significantly reduced near TSS of down-regulated DEGs upon *Ipmk*KD compared to *Egfp*KD (Figure 6D). We further confirmed this by monitoring the specific down-regulated DEG loci, including *Phactr3*, *Lrrc61*, and *Arhgap44* (Figure 6B, right, E, and F). Together, these results suggest that IPMK maintains the expression of a subset of genes by safeguarding the appropriate BRG1 occupancy and BRG1-mediated chromatin accessibility at promoter-TSS in mESCs.

**Figure 6.**
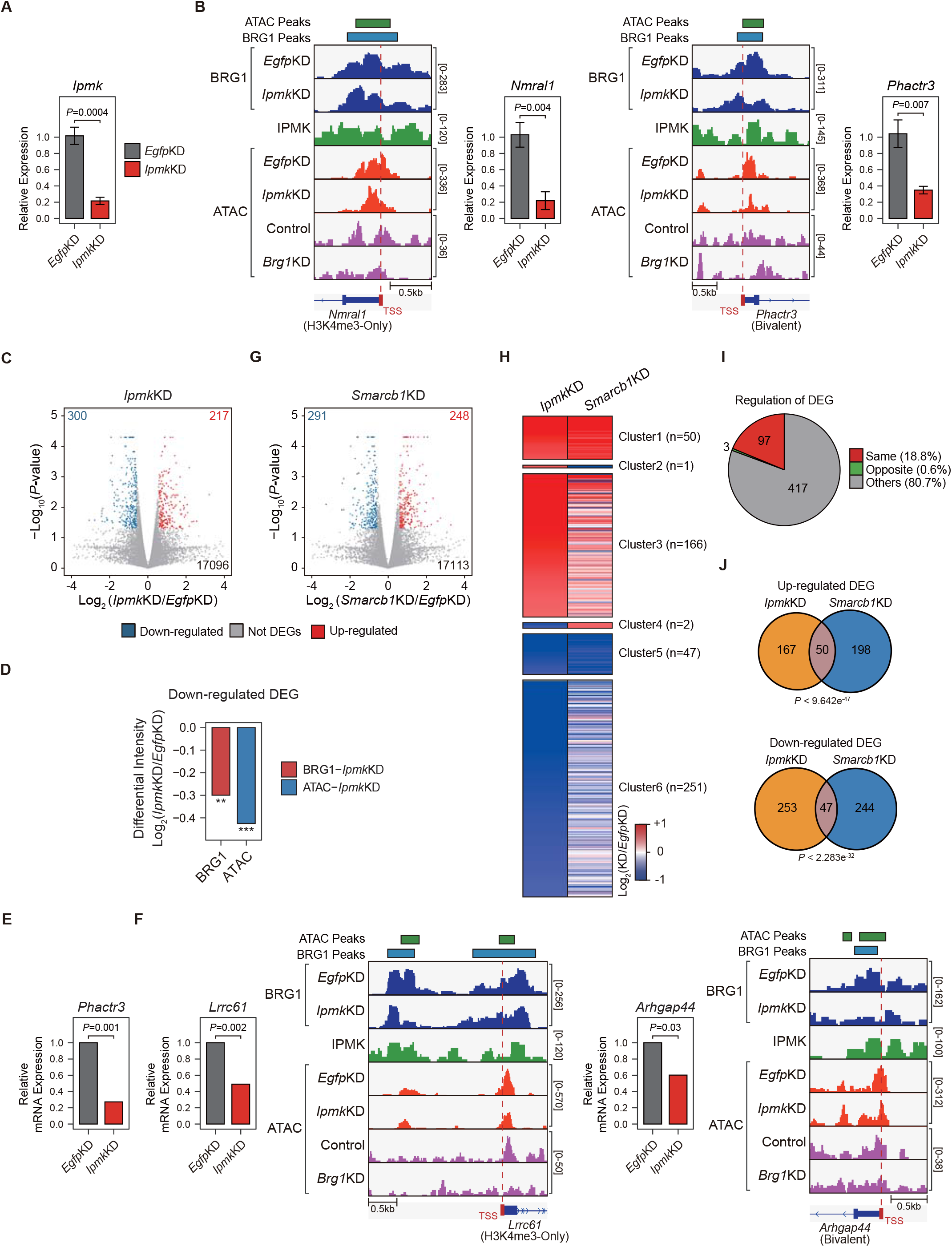
Alteration in BRG1/ATAC upon *Ipmk*KD affects gene expression, and IPMK-SMARCB1 regulates a common set of genes. (A) RT-qPCR analysis of *Ipmk* expression after siRNA treatment. Error bars denote the standard deviation obtained from four biological replicates. The expression levels were normalized with respect to that of β-actin. *P*-value was calculated using Student’s t-test. (B) Examples of BRG1 (*Egfp*KD and *Ipmk*KD), IPMK CUT&RUN, and ATAC-seq (*Egfp*KD, *Ipmk*KD, Control, and *Brg1*KD) assays at TSS of *Nmral1* (left) and *Phactr3* (right). The BRG1 CUT&RUN peaks (*Egfp*KD cells) and ATAC-seq peaks are marked as blue and green boxes on top. TSS are marked with red boxes (bottom) and dotted lines. RT-qPCR analysis of *Nmral1* (left) and *Phactr3* (right) expression after siRNA treatment. Error bars denote the standard deviation obtained from four biological replicates. The expression levels were normalized with respect to that of β-actin. *P*-value was calculated using Student’s t-test. (C) Volcano plots showing the differentially expressed genes (DEGs) upon *Ipmk*KD, identified based on mRNA-seq data. Red and blue dots indicate DEGs that were found to be significantly up-and down-regulated, respectively (*P*-value ≤ 0.05 and fold change ≥ 1.5). (D) Bar graphs showing the average of differential BRG1 (red) and ATAC-seq (blue) intensity upon *Ipmk*KD at BRG1 CUT&RUN peaks (for BRG1 intensity) and at ATAC-seq peaks (for ATAC intensity) that are closest (within 2kb for BRG1 peaks and within 500bp for ATAC-seq peaks) to the TSS of down-regulated DEGs. *P*-values were derived using Wilcoxon signed rank test (**P < 1×10^-5^; ***P < 1×10^-10^). (E) mRNA-seq analysis of *Phactr3* expression after siRNA treatment. *P*-value was calculated using Student’s t-test. (F) Examples of BRG1 (*Egfp*KD and *Ipmk*KD), IPMK CUT&RUN, and ATAC-seq (*Egfp*KD, *Ipmk*KD, Control, and *Brg1*KD) assays at TSS of *Lrrc61* (left) and *Arhgap44* (right). The BRG1 CUT&RUN peaks (*Egfp*KD cells) and ATAC-seq peaks are marked as blue and green boxes on top. TSS are marked with red boxes (bottom) and dotted lines. mRNA-seq analysis of *Lrrc61* (left) and *Arhgap44* (right) expression after siRNA treatment. *P*-value was calculated using Student’s t-test. (G) Volcano plots showing the differentially expressed genes (DEGs) upon *Smarcb1*KD, identified based on mRNA-seq data. Red and blue dots indicate DEGs that were found to be significantly up-and down-regulated, respectively (*P*-value ≤ 0.05 and fold change ≥ 1.5). (H) Heatmaps representing differential gene expression (log_2_ (KD/*Egfp*KD)) of up-regulated (clusters1, 2, and 3) and down-regulated (clusters 4, 5, and 6) DEGs upon *Ipmk*KD. The heatmaps are classified into six clusters based on the differential gene expression of *Ipmk*KD and *Smarcb1*KD cells. (I) Pie chart showing the proportion of *Ipmk*KD-induced DEGs that are regulated in the same manner (red, clusters 1 and 5) or the opposite manner (green, clusters 2 and 4) upon *Ipmk*KD and *Smarcb1*KD. (J) Venn diagrams representing up-regulated DEGs (top) and down-regulated DEGs (bottom) in *Ipmk*KD (orange) and *Smarcb1*KD (blue) cells. The *P*-values indicate the significance of the overlap between the two groups.

### IPMK and SMARCB1 regulate a common set of genes in mESCs and MEFs

To investigate the functional interactions between IPMK and SMARCB1 in a transcription aspect, we performed mRNA-seq using mESCs upon *Ipmk*KD or *Smarcb1*KD. Using the same criteria (fold change ≥ 1.5 and *P*-value ≤ 0.05), we defined differentially expressed genes (DEGs) by comparing the gene expression of *Ipmk*KD or *Smarcb1*KD cells with that of control (*Egfp*KD) cells. A total of 217 up-regulated DEGs and 300 down-regulated DEGs were identified in *Ipmk*KD cells (Figure 6C), and 248 up-regulated DEGs and 291 down-regulated DEGs were identified in *Smarcb1*KD cells (Figure 6G). Given that our previous results demonstrated the physical binding between IPMK and SMARCB1, we hypothesized that IPMK and SMARCB1 would functionally interact with each other. To check this hypothesis, we classified DEGs of *Ipmk*KD cells into six clusters based on the combinatorial mRNA expression changes upon *Ipmk*KD and *Smarcb1*KD (Figure 6H and Supplementary Table 3). Among these six clusters, Cluster1/2/3 represent the up-regulated DEGs upon *Ipmk*KD, whereas Cluster4/5/6 represent the down-regulated DEGs upon *Ipmk*KD. Specifically, Cluster1 and Cluster5 each represent the up-regulated DEGs and down-regulated DEGs shared by both *Ipmk*KD and *Smarcb1*KD cells, respectively. In contrast, Cluster2 and Cluster4 each represent the oppositely regulated DEGs upon *Ipmk*KD and *Smarcb1*KD; Cluster2 contains up-regulated DEG upon *Ipmk*KD and down-regulated DEG upon *Smarcb1*KD, while Cluster4 contains down-regulated DEGs upon *Ipmk*KD and up-regulated DEG upon *Smarcb1*KD. The remaining DEGs of *Ipmk*KD cells were marked as Cluster3/6. Surprisingly, DEGs that were regulated in the same manner (both up-regulated Cluster1 and both down-regulated Cluster5, 18.8%) were far more abundant than DEGs that were oppositely regulated (Cluster2 and Cluster4, 0.6%) (Figure 6H and I). If we assume that IPMK is not functionally related to SMARCB1, the number of DEGs regulated in the same fashion or the opposite fashion should be similar. However, we observed that DEGs regulated in the same manner are much more abundant (∼32 fold more) than DEGs that are oppositely regulated upon *Ipmk*/*Smarcb1*KD, indicating that IPMK and SMARCB1 regulate the common gene sets in mESCs. In addition, we observed that both up-regulated and down-regulated DEGs of *Ipmk*KD cells significantly overlapped with those of *Smarcb1*KD cells (Figure 6J), further supporting the functional linkage between IPMK and SMARCB1. Finally, we performed gene ontology (GO) analysis using genes representing Cluster1 and Cluster5. Despite the similar number of genes in Cluster1 (n=50) and Cluster5 (n=47), the GO terms of Cluster5 were significantly enriched compared to those of Cluster1 (Figure 6—figure supplement 1A), suggesting that down-regulated DEGs regulated by IPMK/SMARCB1 are more closely associated/related to each other compared to up-regulated DEGs. These results indicate that IPMK and SMARCB1 regulate the expression of common gene sets in the same manner, and among these genes, down-regulated DEGs are closely associated with each other.

To further investigate whether the functional interaction between IPMK and SMARCB1 is a specific feature of mESCs or a more general feature, we performed identical analyses in MEFs (NIH3T3). We first defined DEGs; a total of 209 up-regulated DEGs and 275 down-regulated DEGs were identified upon *Ipmk*KD, and 404 up-regulated DEGs and 678 down-regulated DEGs were identified upon *Smarcb1*KD (Figure 6—figure supplement 1B). We then defined the six clusters with the same procedure described for the mESCs. Consistent with the results obtained in the mESCs (Figure 6H and I), DEGs regulated in the same manner (both up-regulated Cluster1 and both down-regulated Cluster5, 23.8%) were much more abundant than DEGs that are oppositely regulated (Cluster 2/4, 1.7%) (Figure 6—figure supplement 1C, D, and Supplementary Table 4). Furthermore, we confirmed that both up-regulated and down-regulated DEGs of *Ipmk*KD cells significantly overlapped with those of *Smarcb1*KD cells (Figure 6—figure supplement 1E), consistent with the results obtained from mESCs (Figure 6J). Taken together, the fact that IPMK and SMARCB1 regulate a common set of genes in the same fashion strongly implies the transcription-related functional interaction between IPMK and SMARCB1 both in mESCs and MEFs.

## DISCUSSION

Using two unbiased screening (yeast two-hybrid and APEX2 proximity labeling), we detected SMARCB1 and other core subunits of the mammalian SWI/SNF complex (BRG1, BAF155, BAF170, ARID1A, and PBRM1) as IPMK-proximal/binding targets. Notably, our binary protein interaction assays showed that IPMK directly binds to SMARCB1, BRG1, and BAF155, individually. Furthermore, *in vivo* and *in vitro* immunoprecipitation assays confirmed that IPMK physically associates with the native SWI/SNF complexes. Detailed mapping studies further revealed reciprocal interactions between the Rpt domains of SMARCB1 and the IP kinase domain of IPMK. In accordance with our previous finding (physical association of IPMK-SWI/SNF complex), our CUT&RUN analysis showed that IPMK is co-localized with BRG1 globally, especially at the promoter-TSS. Surprisingly, by performing CUT&RUN and ATAC-seq, we discovered that the depletion of IPMK severely perturbed (decreased) the genome-wide BRG1 localizations and corresponding BRG1-mediated chromatin accessibility (ATAC-seq signals). By categorizing promoter-TSS, we found that IPMK depletion significantly affected the chromatin accessibility at the bivalent promoters, which were associated with the disrupted BRG1 occupancy at the −1 nucleosome. Using RT-qPCR and mRNA-seq, we detected that the IPMK loss also affects the transcription of genes exhibiting disrupted BRG1/ATAC-seq levels at their promoter-TSS. Lastly, our mRNA-seq analyses also demonstrated that IPMK and SMARCB1 regulate common gene sets, implying a functional link between IPMK and SWI/SNF complex. Taken together, these results elucidate the critical role of IPMK in regulating BRG1 localizations and BRG1-mediated chromatin accessibility through the physical association between IPMK and SWI/SNF complex (Figure 7).

**Figure 7.**
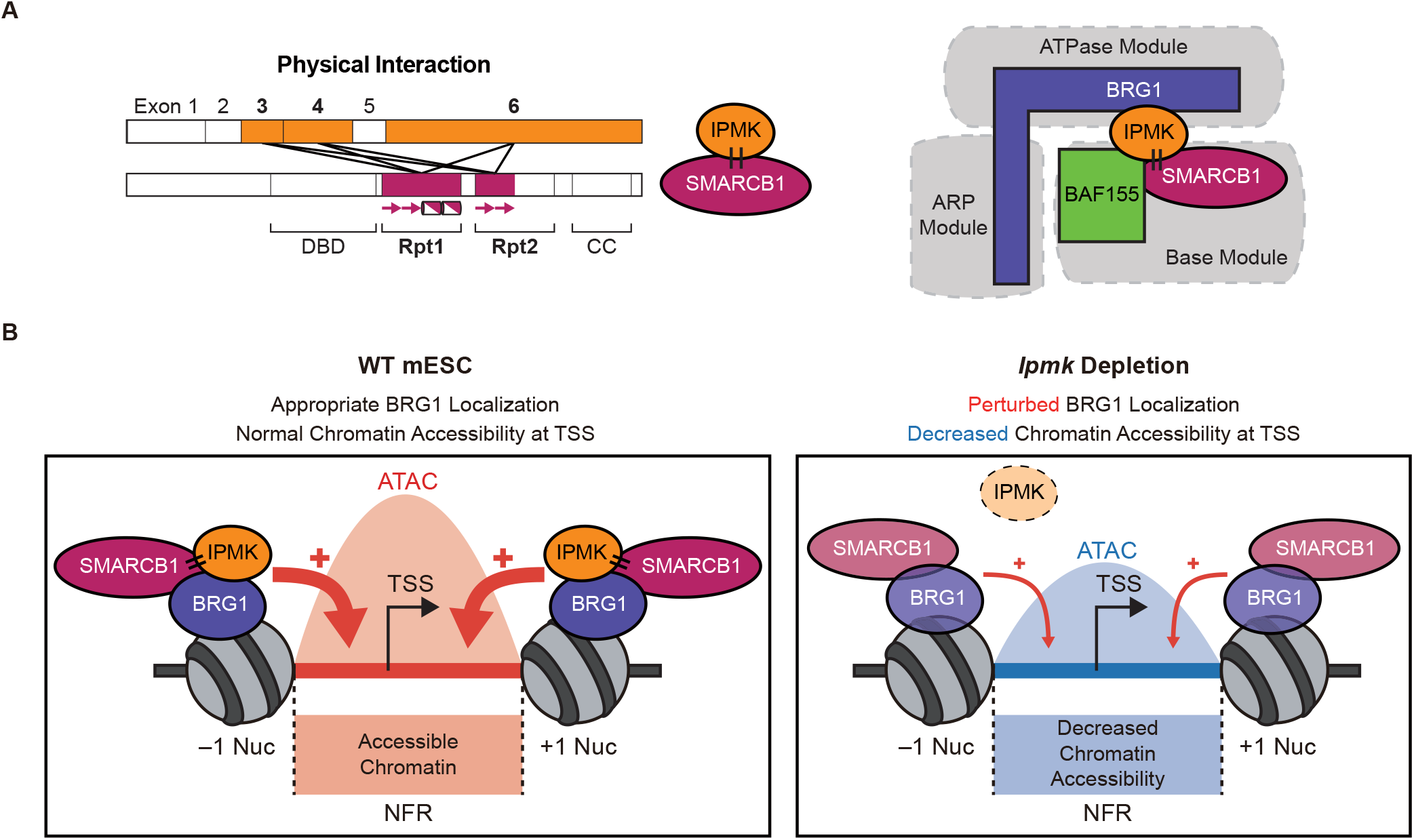
Proposed model depicting the function of IPMK. (A) A model displaying the physical interactions between IPMK and SMARCB1 (left). For these physical interactions, the exons 3, 4, and 6 of IPMK (orange boxes) and the Rpt1 and Rpt2 (particularly N-terminal β sheets) domains of SMARCB1 (red boxes) are required. An additional model showing our speculation on position of IPMK within the SWI/SNF complex, directly interacting with SMARCB1, BRG1, and BAF155 (right). (B) In WT mESC (left), IPMK regulates appropriate BRG1 localization (probably via physical interaction with various subunits of SWI/SNF complex) and chromatin accessibility at NFR (nucleosome free region) of TSS. Upon *Ipmk* depletion (right), BRG1 localization is perturbed, resulting in decreased chromatin accessibility at NFR of TSS.

The physical association between IPMK and SWI/SNF complex is very engaging regarding the structural aspect. According to the recently published cryo-EM-based structure of nucleosome-bound human SWI/SNF complexes (He et al., 2020), the nucleosome is sandwiched by the ATPase module of BRG1 and the base module of SMARCB1 (C-terminal α helix of the Rpt2 domain). BRG1 grasps the upper side of a nucleosome, whereas SMARCB1 binds to the bottom surface of a nucleosome through their positively charged four arginine residues at the C-terminal α helix of Rpt2 domain interacting with the acidic patch of histone octamers (He et al., 2020) (Figure 7—figure supplement 1). Considering this, we revisited our domain mapping results, showing that Rpt1 (all regions including N-terminal β sheets and C-terminal α helices) and Rpt2 (N-terminal β sheets) domains of SMARCB1 and reciprocally, the Exon3/4/6 of IPMK (consists of ∼71% of IPMK proteins) are required for IPMK-SMARCB1 binding (Figure 7A, left). The N-terminal β sheets of the Rpt2 domain seem to be quite buried at the frontal view of nucleosome-bound SWI/SNF complex (He et al., 2020). However, if we rotate the structure and see the back view, we could observe that the beginning part (that is close to C-terminal α helices of Rpt1) of N-terminal β sheets of Rpt2 is being exposed, which may allow some space for IPMK binding. More importantly, we also noticed some space near the Rpt1 domain of SMARCB1 and nucleosomal DNA exit sites, which may provide space for IPMK binding. Intriguingly, this space is positioned near BRG1 and BAF155, which can directly bind to IPMK independent of SMARCB1, according to our results. In accordance with this, our APEX2 data also support this idea by showing that SMARCB1 (BAF47), BRG1 (SMARCA4), SMARCC2 (BAF170), ARID1A (BAF250A), PBRM1 (BAF180), BAF155 (SMARCC1), and core histones (histone H2B, H3.1, and H4) as IPMK-proximal/binding target proteins (Figure 1B, C, and Figure 1—figure supplement 1). Together, we believe that IPMK would be positioned at specific sites within SWI/SNF complex, where it meets the three conditions: (1) sites where IPMK directly binds to the Rpt1 domain (all regions containing N-terminal β sheets and C-terminal α helices) of SMARCB1, (2) sites where IPMK directly binds to SMARCB1, BRG1, and BAF155, and (3) sites proximal to the various subunits of SWI/SNF complexes, including SMARCB1, BRG1, BAF155, BAF170, ARID1A (BAF250A), PBRM1 (BAF180), and core histones. Regarding these conditions, we propose a model in which IPMK physically associates with SWI/SNF complex by directly binding to SMARCB1, BRG1, and BAF155 (Figure 7A, right). Our proposed model also suggests that IPMK may be positioned near the nucleosomal DNA exit site of SWI/SNF complex, which close to the Rpt1 domain of SMARCB1 (Figure 7—figure supplement 1).

It is previously suggested that the Rpt1 domain of SMARCB1 may facilitate the DNA detachment of the SWI/SNF complex, acting as a “wedge” at nucleosomal DNA exit sites (He et al., 2020). According to our results, IPMK is proximal to the core histones and it directly binds to the core subunits (BRG1 and Rpt1 domain of SMARCB1) that sandwich the nucleosomes, suggesting that IPMK may have a role in the interaction between SWI/SNF complex and chromatin. In support of this, our CUT&RUN and ATAC-seq results indicate that the IPMK depletion disrupts the global BRG1 occupancy and its downstream, BRG1-mediated, chromatin accessibility in a genome-wide manner, including promoter-TSS regions. Collectively, our findings suggest that IPMK plays an important role in regulating the localization of mammalian SWI/SNF complex by enhancing the SWI/SNF-nucleosome interactions via physical binding, thereby maintaining the appropriate chromatin accessibility (Figure 7B). Thus, when IPMK is depleted, it affects the SWI/SNF-Nucleosome interactions, resulting in perturbed BRG1 localizations and decreased chromatin accessibility (Figure 7B).

Despite our findings, the mechanism on how IPMK modulates the localization of BRG1 (or SWI/SNF complex) remains unclear. We believe that the physical association of the IPMK-SWI/SNF complex is strongly connected with the BRG1 localization, but the detailed mechanism is still elusive. IPMK may facilitate the recruitment of the SWI/SNF complex’s subunits from cytoplasm to chromatin. However, our observations from chromatin fractionation assays indicated that IPMK depletion did not affect the BRG1 or SMARCB1 occupancy in the cytoplasm or chromatin fractions (Figure 4A), excluding the above possibility. Alternatively, it is plausible that IPMK may aid the appropriate conformation of specific subsets of SWI/SNF complex by physically binding near the nucleosomal DNA exit sites within the SWI/SNF complex. Thus, by aiding the appropriate conformation of the SWI/SNF complex, IPMK plays a vital role in SWI/SNF-nucleosome interactions, thereby facilitating/stabilizing the BRG1 occupancy on the chromatin. Although IPMK depletion reduced the global BRG1 occupancy, our CUT&RUN results indicated that IPMK depletion exhibits more impact at the CUT&RUN peaks with lowly enriched BRG1 in *Egfp*KD (wild-type-like) mESCs, compared to highly enriched BRG1 peaks (Figure 4C and G-I). Importantly, our ATAC-seq results also corresponded to the region-specific (BRG1 low) changes in BRG1 occupancy upon *Ipmk*KD (Figure 4C and G-I). In support of this, IPMK depletion exhibited more impact on the chromatin accessibility and transcription of the relatively-low-BRG1-harboring bivalent promoters than H3K4me3-Only promoters, which harbor high BRG1 levels (Figure 5 and 6). Together, these results indicate that IPMK primarily regulates the BRG1 occupancy (which resembles the SWI/SNF-nucleosome interactions) and its downstream effects (chromatin accessibility and transcription) at the region where the BRG1 level is originally low in mESCs but does not affect the regions with high BRG1 level. This contextual discrepancy in IPMK-dependent or IPMK-independent BRG1 occupancy may provide key clues to the mechanism on how IPMK modulates the BRG1 localization, but further experiments/analyses are required.

Previous studies showing that IPMK regulates several target proteins (including cytosolic signaling factors and transcription factors) through protein-protein interaction also support our view. In earlier studies, IPMK was found to bind to mTOR and raptor, maintaining the mTOR-raptor association and amino acid-induced mTOR signaling (S. Kim et al., 2011). Glucose signaling activates the phosphorylation of IPMK, enabling IPMK-AMPK binding, thereby enhancing its signaling (Bang et al., 2012). Furthermore, IPMK stimulates p53-mediated transcription by binding to p53, thus facilitating p53-mediated cell death (Xu, Sen, et al., 2013). IPMK also binds with CBP/p300, a transcriptional coactivator of CREB, and augments the expression of CREB-regulated genes (Xu, Paul, et al., 2013). IPMK also binds to SRF, enhancing its interactions with the serum response element at the promoters, and induces immediate early gene expressions (E. Kim et al., 2013). Lastly, IPMK, which possesses PI3K activity, interacts with the SF-1 nuclear receptor and generates SF-1–PIP_3_ complex, promoting lipid-mediated signaling in the nucleus (Blind et al., 2014; Blind, Suzawa, & Ingraham, 2012). Conversely, core subunits of the SWI/SNF complex also interact with various factors, such as PDX1 and CTCF (Marino et al., 2019; Spaeth et al., 2019). In addition, direct interactions between SWI/SNF complex and oncogenes/tumor-suppressor genes, such as RB (Retinoblastoma), BRCA1, c-MYC, and MLL, have been implicated in oncogenesis (Roberts & Orkin, 2004). Although these studies support our view that the direct physical association between IPMK and SWI/SNF complex may affect diverse epigenetic events (such as BRG1 localization, chromatin accessibility, or transcription), further experiments are required to elucidate the precise underlying mechanisms of this phenomenon.

SWI/SNF chromatin remodeling complex is frequently mutated in cancer (Helming, Wang, & Roberts, 2014). Particularly, SMARCB1 is a *bona fide* tumor suppressor gene (Roberts, Galusha, McMenamin, Fletcher, & Orkin, 2000; Roberts, Leroux, Fleming, & Orkin, 2002), which is inactivated or lost in multiple malignancies, such as malignant rhabdoid tumors. A germline mutation of *IPMK* was recently found in familial small intestinal carcinoid patients (Sei et al., 2015). A truncated *IPMK* allele was associated with reduced P53 signaling in these patients, suggesting a tumor suppressor role of IPMK. Additionally, the *IPMK* mRNA levels were down-regulated by the tumor suppressor miR-18a, which contributes to the inhibition of ovarian tumor growth (Liu et al., 2017). Finally, the exogenous supply of IP_4_, which is produced by IPMK, can suppress human cancer cell growth by inhibiting the activation of AKT/PKB (Jackson, Al-Saigh, Schultz, & Junop, 2011; Piccolo et al., 2004; Razzini et al., 2000). Considering these previous reports with our results (that elucidated the physical/function linkage between the tumor suppressor SMARCB1 and IPMK), it would be interesting to conduct genome-wide studies characterizing the role of combined action of IPMK-SMARCB1 in the context of cancer.

Our study is the first to elucidate the physical association between IPMK and core subunits of SWI/SNF complex, and the first to define the molecular function of IPMK in coordinating the BRG1 localizations and BRG1-associated chromatin accessibility in mESCs. Considering our results and recently published cryo-EM-based structure of human SWI/SNF (BAF) complex, we propose a model in which IPMK physically associates with SWI/SNF complex via directly binding to SMARCB1, BRG1, and BAF155 and positions near the nucleosomal DNA exit site of SWI/SNF complex, which is in close proximity to the Rpt1 domain of SMARCB1 (the IPMK-binding domain). Based on this model, IPMK plays an important role in regulating the SWI/SNF-nucleosome interactions, thereby maintaining an appropriate BRG1 occupancy and BRG1-mediated chromatin accessibility. We believe these novel findings will play a pivotal role in future studies of IPMK and understanding the molecular mechanisms of mammalian SWI/SNF complexes, especially by providing additional clues in SWI/SNF-mediated generation of nucleosome free regions at the transcription start sites (TSS).

## MATERIALS AND METHODS

### Yeast two-hybrid screening

Panbionet (Pohang, South Korea) conducted yeast two-hybrid screening (http://panbionet.com). The full IPMK coding region of 416 amino acids was amplified by polymerase chain reaction (PCR). The PCR product was cloned into the pGBKT7 vector, which contains the DNA-binding domain (BD) of GAL4. *Saccharomyces cerevisiae* strain AH109 (Clontech) was co-transformed with GAL4 DNA-BD-fused IPMK and a human brain cDNA activation domain (AD) library (Clontech). Two different reporter genes (*HIS3* and *ADE2*) were used as selection markers. Yeast transformants were spread on a selection medium lacking leucine, tryptophan, and adenine or histidine (SD−LWA and SD-LWH). To confirm the interactions, the candidate prey genes of candidates were amplified via PCR or *E. coli* transformation, and reintroduced into the AH109 yeast strain with the IPMK bait plasmid.

### Generation of stable cell lines for APEX2-mediated proximity labeling

V5-APEX2 was PCR-amplified from the pcDNA5-Mito-V5-APEX2 plasmid, which was kindly provided by Dr. Hyun-Woo Rhee (Seoul National University). V5-APEX2 alone or IPMK-V5-APEX2 were cloned into the pcDNA™5/FRT/TO plasmid (Invitrogen). Flp-In™ T-REx™-293 (Invitrogen) cells were seeded in 6-well culture plate to reach 70% confluency, then co-transfected with 0.25 µg of pcDNA™5/FRT/TO and 2.25 µg of pOG44 Flp recombinase expression plasmid (Invitrogen) using Lipofectamine LTX with Plus Reagent (Invitrogen). After 48 hours, the cells were transferred to 90 mm culture dishes to undergo negative selection with 50 µg/ml Hygromycin B (Gibco) until all non-transfected cells were dead. Surviving cells were then seeded with low confluency to generate cellular clones on culture plates, after which each clone was individually screened for APEX2 construct expression with or without doxycycline (Sigma Aldrich) in order to search for optimal cell populations with minimal uncontrolled APEX2 expression and maximal APEX2 expression under stimulation. Selected clones were then expanded and stored in liquid nitrogen for downstream experiments.

### APEX2-mediated proximity labeling

1.4×10^7^ APEX2-expressing cells were seeded in T75 culture flasks. 16 hours after seeding, the culture medium was exchanged with complete medium supplemented with doxycycline (100 ng/ml) for APEX2 expression. After 24 hours of induction, the cells were incubated in fresh media containing 250 µM desthiobiotin-phenol (DBP) for 30 minutes in a CO_2_ incubator at 37°C. The cells were then moved to room temperature, and hydrogen peroxide (diluted in DPBS to 1mM; Sigma Aldrich) was added to initiate the APEX2-driven biotinylation reaction. The reaction was quenched by adding a 2X quenching solution (20 M sodium ascorbate, 10 mM Trolox, and 20 mM sodium azide in DPBS) to the medium. The cells were further washed with 1X quencher solution for three times, collected by centrifugation, snap-frozen and stored at −80°C until lysis. DBP was synthesized as described in a previous report (S. Y. Lee et al., 2017).

### Preparation of DBP-labeled peptides for LC-MS/MS

DBP-labeled peptides were prepared from frozen cell pellets as described in a previous report (Kwak et al., 2020). Briefly, cells were lysed in lysis buffer (2% SDS, 1X protease inhibitor cocktail (Roche), and 1 mM sodium azide in 1X TBS) and excess DBP was eliminated through repeated acetone precipitation. The resulting protein precipitates were again solubilized in 50 mM ammonium bicarbonate and quantified. 4 mg of cellular protein was then denatured, reduced, alkylated, and digested into peptides with trypsin. Afterward, tryptic DBP-labeled peptides were bound to streptavidin beads (Pierce) and collected with elution buffer (80% acetronitrile, 0.2% trifluoroacetic acid, 0.1% formic acid in MS-grade water). Solvents were completely evaporated on a SpeedVac for 3 hours, and the resulting peptides were stored at −20°C until required for LC-MS/MS analysis.

### LC-MS/MS

The resulting tryptic peptides were analyzed by LC-MS/MS. All mass analyses were performed on a Q Exactive Plus orbitrap mass spectrometer (Thermo Fisher Scientific) equipped with a nanoelectrospray ion source. To separate the peptide mixture, we used a C18 reverse-phase HPLC column (500 mm × 75 μm ID) using an acetonitrile/0.1% formic acid gradient from 3.2 to 26% for 120 minutes at a flow rate of 300 nL/min. For MS/MS analysis, the precursor ion scan MS spectra (m/z 400∼2000) were acquired in the Orbitrap at a resolution of 70,000 at m/z 400 with an internal lock mass. The 15 most intensive ions were isolated and fragmented by High-energy collision induced dissociation (HCD).

### LC-MS/MS data processing

All MS/MS samples were analyzed using Sequest Sorcerer platform (Sagen-N Research, San Jose, CA). Sequest was set up to search the *Homo sapiens* protein sequence database (20675 entries, UniProt (http://www.uniprot. org/)), which includes frequently observed contaminants, assuming the digestion enzyme trypsin. Sequest was searched with a fragment ion mass tolerance of 1.00 Da and a parent ion tolerance of 10.0 ppm. Carbamidomethylation of cysteine was specified in Sequest as a fixed modification. Oxidation of methionine and acetyl of the n-terminus, biotin of lysine and DBP of tyrosine were specified in Sequest as variable modifications. Scaffold (Version 4.11.0, Proteome Software Inc., Portland, OR) was used to validate MS/MS-based peptide and protein identifications. Peptide identifications were accepted if they could be established at greater than 93.0% probability to achieve a false discovery rate (FDR) less than 1.0% by the Scaffold Local FDR algorithm. Protein identifications were accepted if they could be established at greater than 92.0% probability to achieve an FDR less than 1.0% and contained at least two identified peptides. Protein probabilities were assigned by the Protein Prophet algorithm (Nesvizhskii, Keller, Kolker, & Aebersold, 2003). Proteins that contained similar peptides and could not be differentiated based on MS/MS analysis alone were grouped to satisfy the principles of parsimony. Proteins were annotated with GO terms from NCBI (downloaded November 23, 2019) (Ashburner et al., 2000).

### Plasmids

The cDNAs for human IPMK (NCBI Gene ID 253430) and human SMARCB1 (NCBI Gene ID 6598) were obtained respectively from Open Biosystems and Bioneer (Daejeon, South Korea). IPMK and SMARCB1 cDNA constructs were amplified by PCR and the products were cloned into pCMV-GST and pcDNA3.1-FLAG vectors. Every construct was confirmed by DNA sequencing.

### *In vitro* binding assay

Recombinant human IPMK was purified as described previously (B. Lee et al., 2020). Briefly, human IPMK was expressed in Sf9 insect cells with a baculovirus system, harvested with lysis buffer consisting of 300 mM NaCl, 50 mM Tris, pH 8.0, 5% glycerol and 1 mM phenylmethylsulfonylfluoride (PMSF). Freezing and thawing lysis method with liquid nitrogen was applied to the cells, and the supernatants were taken after centrifugation at 18,000 rpm for 90 minutes. Ni-NTA agarose (Qiagen) was applied with 20 mM imidazole incubated for 2 hours. The protein was eluted with 100 mM imidazole and the N-terminal HIS-tag was removed with TEV protease, followed by further purification with HiTrap and Superdex columns (GE Healthcare). Human SMARCB1 was translated *in vitro* using TNT Quick Coupled Transcription/Translation System (L1170, Promega). 1 µg of pcDNA3.1-FLAG-SMARCB1 was incubated at 30°C for 90 minutes with 20 µM methionine and TNT T7 Quick Master Mix. Translated FLAG-SMARCB1 was incubated with anti-FLAG M2 affinity gel (A2220, Sigma Aldrich), and then IPMK protein was added and incubated with rotation at 4°C.

### Recombinant IPMK protein purification

For GST-tagged protein, human IPMK cDNA was subcloned into pGEX4T plasmid (Sigma Aldrich), expressed in Escherichia coli, and purified on Glutathione Sepharose 4B beads (GE Healthcare) as described (J. Kim & Roeder, 2011). For FLAG-tagged proteins, wild-type or mutant IPMK cDNAs were subcloned into pFASTBAC1 plasmid (Thermo Fisher Scientific) with an N-terminal FLAG epitope, and baculoviruses were generated according to the manufacturer’s instructions. Proteins were expressed in Sf9 insect cells and purified on M2 agarose (Sigma Aldrich) as described (J. Kim & Roeder, 2011).

### BAF complex purification

The FLAG-DPF2 cell line was selected from HEK293T cells transfected with a FLAG-DPF2-pCAG-IP plasmid. Derived nuclear extracts (Dignam, Lebovitz, & Roeder, 1983) were incubated with M2 agarose in binding buffer (20 mM Tris HCl [pH 7.3], 300 mM KCl, 0.2 mM EDTA, 25% glycerol, 1.5 mM MgCl_2_, 10 mM 2-mercaptoethanol. and 0.2 mM PMSF) at 4°C for 4 h. After extensive washing with wash buffer (20 mM Tris HCl [pH 7.9], 150 mM NaCl, 0.2 mM EDTA, 5% glycerol, 2 mM MgCl_2_, 10 mM 2-mercaptoethanol, 0.2 mM PMSF, and 0.1% NP-40), complexes were eluted with wash buffer containing 0.25 mg/ml FLAG peptide. Eluted complexes were fractionated by 10%–30% glycerol gradient and the fractions containing intact BAF complex were combined and concentrated using Amicon Ultra-4 centrifugal filter (Millipore).

### Protein interaction assays

For GST pull-down assays, 2ug of GST or GST-tagged IPMK immobilized on Glutathione Sepharose 4B beads were incubated with 200 ng of purified BAF complexes in binding buffer (20 mM Tris-HCl [pH 7.9], 150 mM KCl, 0.2 mM EDTA, 20% glycerol, 0.05% NP-40, and 0.2 mg/ml BSA) at 4°C for 3 h. Beads were extensively washed with binding buffer without BSA, and bound proteins were analyzed by immunoblotting. For binary protein interaction assays following baculovirus-mediated expression, Sf9 cells were infected with baculoviruses expressing FLAG-IPMK and untagged BAF complex subunit. After 2 days, total cell extracts were prepared by sonication in lysis buffer (20 mM Tris-HCl [pH7.9], 300 mM NaCl, 0.2 mM EDTA, 15% glycerol, 2 mM MgCl_2_, 1 mM DTT, 1 mM PMSF, and protease inhibitor cocktail [Roche]). Following clarification by centrifugation, cell extracts were incubated with M2 agarose at 4°C for 3 h and, after extensive washing with wash buffer (20 mM Tris-HCl [pH 7.9], 150 mM NaCl, 0.2 mM EDTA, 15% glycerol, 2 mM MgCl_2_, 1 mM DTT, 1 mM PMSF, and 0.1% NP-40), bound proteins were analyzed by immunoblotting.

### Immunoblotting, immunoprecipitation, and GST pull-down

For immunoblot analyses, the cells were washed twice with PBS and lysed in lysis buffer consisting of 1% Triton X-100, 120 mM NaCl, 40 mM Tris-HCl, pH 7.4, 1.5 mM sodium orthovanadate, 50 mM sodium fluoride, 10 mM sodium pyrophosphate, 1 mM EDTA, and protease inhibitor cocktail (Roche). Cell lysates were incubated at 4°C for 10 minutes, and the supernatants were collected by centrifuging at 13,000 rpm for 10 min. Protein concentrations were determined by the Bradford protein assay (Bio-Rad) or bicinchoninic acid (BCA) assay (Thermo Fisher Scientific). 20 µg of protein lysates were separated by size, transferred to nitrocellulose membranes, and blotted with primary antibodies and secondary antibodies. The horseradish peroxidase (HRP) signals were visualized with the Clarity ECL substrate (Bio-Rad) and SuperSignal^TM^ West Femto Maximum Sensitivity Substrate (Thermo Fisher Scientific), and measured by using a ChemiDoc imaging system (Bio-Rad). For immunoprecipitation, 2 mg of total protein was incubated with 5 µg of primary antibodies for 16 hours with rotation at 4°C. 10 µL of TrueBlot beads (Rockland Immunochemicals) were added and incubated for an additional hour. The samples were washed three times with lysis buffer and prepared for immunoblotting. For GST pull-down assay, 10 µL of glutathione agarose beads (Incospharm) were added to 2 mg of total cell lysates and incubated for 16 hours with rotation at 4°C. The samples were then washed three times with lysis buffer and prepared for immunoblotting.

### Cell culture and cell line production

E14Tg2a mouse embryonic stem cells (mESCs) were maintained under feeder-free conditions. Briefly, the cells were cultured on gelatin-coated cell culture dishes in an mESCs culture medium consisting of Glasgow’s minimum essential medium (GMEM) containing 10% knockout serum replacement, 1% non-essential amino acids, 1% sodium pyruvate, 0.1 mM β-mercaptoethanol (all from Gibco), 1% FBS, 0.5% antibiotic-antimycotic (both from Hyclone) and 1,000 units/ml LIF (ESG1106, Millipore). The mESCs were maintained at 37°C with 5% CO_2_ in humidified air. NIH3T3 cells, mouse embryonic fibroblasts (MEFs) and human embryonic kidney (HEK)-293T cells were grown in high-glucose DMEM supplemented with 10% FBS, 2 mM L-glutamine, and penicillin/streptomycin (100 mg/ml), and maintained in a humid atmosphere of 5% CO_2_ at 37°C. To generate tamoxifen-inducible IPMK knockout mice, *Ipmk^fl/fl^* mice were mated with UBC-Cre-ERT2 mice (The Jackson Laboratory). The MEFs were immortalized by transfecting with an SV40 large T-antigen plasmid, and the IPMK depletion was achieved by adding 1 µM 4-hydroxytamoxifen for 48 hours. FLAG epitope-tagged mESCs and MEFs were generated as described previously (Savic et al., 2015). Briefly, the 3xFlag-P2A-Puromycin epitope tagging donor construct (pFETCh-Donor), CRISPR guide RNAs (gRNAs), and Cas9 expressing plasmids were manufactured by ToolGen (Seoul, Korea). mESCs were transfected using FUGENE HD (E2311, Promega), selected using puromycin (A11138-03, Gibco), and expanded. MEFs were transfected with the donor construct containing the neomycin resistance gene with Turbofect (R0533, Thermo Fisher Scientific) and selected using G418 (11811023, Gibco).

### RNA interference

Control siRNA (scRNA) and siRNAs against *Egfp* (sense: 5’-GUUCAGCGUGUCCGGCGAG-3’; antisense: 5’-CUCGCCGGACACGCUGAAC-3’) and *Ipmk* (5′-CAGAGAGGUCCUAGUUAAUUUCA-3′; antisense: 5′-AGUGAAAUUAACUAGGACCUCUCUGUU-3′) were synthesized and annealed by Bioneer (Daejeon, Korea), and siRNA against *Smarcb1* was purchased from Sigma Aldrich. mESCs and MEFs were transfected with 50 nM of the corresponding siRNA using DharmaFECT I (T-2001-03, Dharmacon) according to the manufacturer’s instructions. Briefly, the cells were seeded onto 6-well plates. One day later, 50 nM of siRNAs and DharmaFECT reagent were diluted in Opti-MEM (Gibco), incubated separately at 25°C for 5 minutes, and then mixed together. The mixtures were incubated at 25°C for 20 minutes and added to the cell cultures. The culture medium was replaced after 24 hours and the transfected cells were harvested at 48 hours after transfection.

### Chromatin fractionation

A chromatin acid extraction was performed as described previously (Zhong, Martinez-Pastor, Silberman, Sebastian, & Mostoslavsky, 2013). mESCs or MEFs were collected and washed with PBS, and resuspended with lysis buffer consisting of 10 mM HEPES, pH 7.4, 10 mM KCl, 0.05% NP-40, 1 mM sodium orthovanadate, and protease inhibitor cocktail (Roche). The cell lysates were incubated 20 minutes on ice and centrifuged at 13,000 rpm. The supernatant contained the cytoplasmic proteins, and the pellet with nuclei was washed once with lysis buffer and centrifuged at 13,000 rpm for 10 minutes. The nuclei were resuspended with low salt buffer consisting of 10 mM Tris-HCl, pH 7.4, 0.2 mM MgCl_2_, 1% Triton-X 100, 1 mM sodium orthovanadate, and protease inhibitor cocktail, then incubated 15 minutes on ice. After 10 minutes of centrifugation, the supernatant contained the nucleoplasmic proteins, and the pellet contained the chromatin. The chromatin was then resuspended with 0.2 N HCl for 20 minutes on ice, centrifuged at 13,000 rpm for 10 minutes, and neutralized with 1M Tris-HCl, pH 8.0. The protein concentrations were determined and subjected to immunoblotting.

### CUT&RUN

CUT&RUN assays were performed as previously described (Meers, Bryson, Henikoff, & Henikoff, 2019; Skene & Henikoff, 2017), with minor modification. Briefly, 4 million mESCs were harvested and washed with 1.5 ml Wash buffer (20 mM HEPES, pH 7.5, 150 mM NaCl, 0.5 mM Spermidine) three times. Cells were bound to activated Concanavalin A-coated magnetic beads (at 25°C for 10 min on a nutator), then permeabilized with Antibody buffer (Wash buffer containing 0.05% Digitonin and 4 mM EDTA). The bead-cell slurry was incubated with 3 μl relevant antibody (see below) in a 150 μl volume at 25°C for 2 hours on a nutator. After two washes in 1 ml Dig-wash buffer (Wash buffer containing 0.05% Digitonin), beads were resuspended in 150 μl pAG/MNase and incubated at 4°C for 1 hour on a nutator. After two washes in 1 ml Dig-wash buffer, beads were gently vortexed with 100 μl Dig-wash buffer. Tubes were chilled to 0°C for 5min and ice-cold 2.2 mM CaCl2 was added while gently vortexing. Tubes were immediately placed on ice and incubated at 4°C for 1 hour on a nutator, followed by addition of 100 μl 2xSTOP buffer (340 mM NaCl, 20 mM EDTA, 4 mM EGTA, 0.05% Digitonin, 0.1 mg/ml RNase A, 50 μg/ml glycogen) and incubated at 37°C for 30min on a nutator. Beads were placed on a magnet stand and the liquid was removed to a fresh tube, followed by addition of 2 μl 10% SDS and 2.5 μl proteinase K (20mg/ml) and incubated at 50°C for 1 hour. DNA was extracted using phenol chloroform as described at https://www.protocols.io/view/cut-amp-run-targeted-in-situ-genome-wide-profiling-zcpf2vn. CUT&RUN libraries were prepared using a NEXTflex ChIP-seq Library kit (5143-02, Bioo Scientific), according to the manufacturer’s guidelines. The libraries were then sequenced using an Illumina Novaseq 6000 platform. The libraries were generated from two sets of biological replicates.

### Antibodies

Antibodies against FLAG (F1804, Sigma Aldrich), IPMK (custom rabbit polyclonal antibody, raised against a mouse IPMK peptide corresponding to amino acids 295-311 (SKAYSTHTKLYAKKHQS; Covance)) (S. Kim et al., 2011), SMARCB1 (A301-087, Bethyl), BRG1 (ab110641, Abcam), BAF155 (11956, Cell Signaling Technology), BAF170 (12760, Cell Signaling Technology), BAF250A (12354, Cell Signaling Technology), BRM (11966, Cell Signaling Technology), PBAF/PBRM (A301-591A, Bethyl), a-TUBULIN (T5169, Sigma Aldrich), GST (2622, Cell Signaling Technology), LaminB1 (sc-365214, Santa Cruz Biotech), Histone H3 (homemade), GAPDH (sc-32233, Santa Cruz Biotech), anti-DPF2 (ab128149, Abcam), anti-SMARCE1 (ab137081, Abcam), anti-SS18L1 (ab227535, Abcam), anti-ACTL6A (sc-137062, Santa Cruz Biotech), anti-SMARCD1 (sc-135843, Santa Cruz Biotech), anti-BCL7A (HPA019762, Atlas Antibodies) and anti-ACTB (TA811000, Origene) were used for immunoblotting. Antibodies against IPMK (homemade), SMARCB1 (A301-087, Bethyl) and normal rabbit IgG (sc-2027, Santa Cruz Biotech) and anti-FLAG M2 affinity gel (A2220, Sigma Aldrich) were used for immunoprecipitation, and GST (2622, Cell Signaling Technology) was used for pulldown. Antibodies against FLAG (F7425, Sigma Aldrich), BRG1 (ab110641, Abcam), and IgG (homemade) were used for CUT&RUN assay.

### ATAC-seq

ATAC-seq libraries were prepared as previously described (Buenrostro, Giresi, Zaba, Chang, & Greenleaf, 2013; Buenrostro, Wu, Chang, & Greenleaf, 2015), with minor modification. Briefly, 50,000 mESCs were harvested, washed with cold PBS, lysed with cold lysis buffer, and immediately centrifuged. The nuclear pellets were resuspended in 25 μl of 2X tagmentation reaction buffer (10 mM Tris, pH 8.0, 5 mM MgCl_2_, 10% dimethylformamide), 23 μl of nuclease-free water, and 2 μl of Tn5 transposase (in-house generated), and incubated at 37°C for 30 min. The samples were then immediately purified using a QIAquick PCR purification kit (28106, Qiagen). The libraries were pre-enriched for five cycles using the KAPA HiFi Hotstart ready mix (KK2601, Kapa Biosystems), and the threshold cycle (Ct) was monitored using qPCR to determine the additional enrichment cycles, which were then applied. The final libraries were purified again with a QIAquick PCR purification kit, and sequenced using an Illumina Novaseq 6000 platform. The libraries were generated from two sets of biological replicates.

### H3K4me3-Only and bivalent promoter-TSS

The list of 47,382 mouse genes was obtained from the UCSC genome browser (Table browser, mm10, group: Genes and Gene Prediction, track: NCBI RefSeq, table: UCSC RefSeq (refGene), region: genome). Among these genes, we selected protein-coding genes (gene name starting with NM_) that are longer than 2 kb. To classify the promoter-TSS regions precisely, we removed redundancies by merging the genes, having the exact same transcription start sites (TSS), into the same group. By doing so, we obtained 23,927 mouse promoter-TSS regions. To categorize the promoter-TSS regions depending on their histone modifications status, we first calculated the H3K4me3 (accession number: GSM254000 and GSM254001) and H3K27me3 (accession number: GSM254004 and GSM254005) ChIP-seq intensity in a −500/+1,000 bp window around the promoter-TSS regions. The −500/+1,000 bp window range was also applied in a previous study (de Dieuleveult et al., 2016). Next, we divided the promoter-TSS into two groups based on the first quartile (Q1) value of H3K4me3 ChIP-seq intensity at the whole promoter-TSS regions; H3K4me3-Low and H3K4me3-High. We then divided the H3K4me3-High promoter-TSS into two groups based on the third quartile (Q3) value of H3K27me3 ChIP-seq intensity at the whole promoter-TSS regions; H3K4me3-Only and bivalent. Thus, we categorized three types of promoter-TSS; 5,982 H3K4me3-Low, 12,305 H3K4me3-Only (high H3K4me3 and low H3K27me3), and 5,640 bivalent (high H3K4me3 and high H3K27me3). We obtained similar results when using the publically released H3K4me3 and H3K27me3 ChIP-seq data (ENCODE, Ross Hardison, ENCSR212KGS and ENCSR059MBO). Furthermore, we confirmed that the percentage of three promoter-TSS types are similar to the results obtained by a previous study (de Dieuleveult et al., 2016).

### mRNA purification and mRNA-seq

Total RNA was purified from mESCs using the TRIzol reagent (Invitrogen) according to the manufacturer’s instructions. Briefly, mESCs cultured in 6-well plates were harvested and homogenized with 1 ml of TRIzol reagent. Chloroform (200 μL/sample) was then added, and the samples were vigorously mixed by hand for 15 seconds and incubated at 25°C for 2 minutes. The mixtures were centrifuged at 12,000 rpm for 15 minutes at 4°C, and 500 μL of each aqueous phase was transferred to a new Eppendorf tube and mixed with the equal volumes of isopropanol. The mixtures were incubated at 25°C for 10 minutes to precipitate the total RNA samples. The samples were then centrifuged at 12,000 rpm for 10 minutes at 4°C, washed with 75% ethanol, and centrifuged again at 10,000 rpm for 5 minutes at 4°C. The RNA pellets were dried and dissolved in RNase-free water. For mRNA sequencing (mRNA-seq) library preparation, mRNA was isolated from total RNA using a Magnetic mRNA isolation kit (S1550S, NEB), and libraries were prepared using a NEXTflex Rapid directional RNA-seq kit (5138-08, Bioo Scientific). The libraries were sequenced using an Illumina HiSeq 2500 system. The libraries were generated from two sets of biological replicates.

### Gene Ontology analysis

ConsensusPathDB was used to identify gene ontology terms associated with differentially expressed genes (DEGs) (Herwig et al., 2016).

### Data processing and analysis

For CUT&RUN analysis, raw reads were aligned to the mouse genome (mm10) using Bowtie2 (version 2.2.9) (Langmead & Salzberg, 2012) with the parameter (--trim3 125 --local --very-sensitive-local --no-unal --no-mixed --no-discordant -q --phred33 -I 10 -X 700), as the previous study (GSM2247138). For ATAC-seq analysis, raw reads were aligned to the mouse genome (mm10) using Bowtie2 (version 2.2.9) with the following parameter; --very-sensitive -X 100 –local. For mRNA-seq analysis, raw reads were aligned to the mouse genome (mm10) using STAR (version 2.5.2a) with default parameters (Dobin et al., 2013). Generally, we used MACS2 (Zhang et al., 2008) to convert the aligned BAM files into bedGraph files, and normalized the data with respect to the total read counts. Then, we used bedGraphToBigWig (Kent, Zweig, Barber, Hinrichs, & Karolchik, 2010) to convert the bedGraph files into bigWig files. The bigWig files were used as input files for bwtool (Pohl & Beato, 2014) (matrix and aggregate option) to quantify the intensity (e.g., heatmaps or average line plots) of the relevant sequencing data. All of our raw data (fastq files) were confirmed to be of good quality using FastQC (http://www.bioinformatics.babraham.ac.uk/projects/fastqc/). For our CUT&RUN analysis, we used MACS2 (callpeak option, P-value < 0.005) to identify the peaks (or binding sites) of protein of interest by using IgG as backgrounds. The CUT&RUN data was also subjected to HOMER annotatPeaks.pl (Heinz et al., 2010) to elucidate the genomic contents within BRG1/IPMK-binding sites. For our mRNA-seq analyses, we used Cufflinks (Trapnell et al., 2010) (Cuffdiff option, fr-firststrand) to assess the expression levels and identify DEGs. Box plots, volcano plots, and other plots were drawn with R (ggplot2) (Wickham, 2009) and heatmaps were drawn with Java TreeView (Saldanha, 2004). The examples of our genome-wide data were visualized using the Integrative Genomics Viewer (IGV) (Robinson et al., 2011).

### Public data acquisition

Publicly released ChIP-seq data were downloaded from the NCBI GEO DataSets database. These data were downloaded as sra or fastq files, and sra files were converted to fastq files using the SRA Toolkit (https://trace.ncbi.nlm.nih.gov/Traces/sra/sra.cgi?view=software). Both the public datasets and our data were then analyzed using the same methods.

## DATA AVAILABILITY

The NGS data from this study have been submitted to the NCBI Gene Expression Omnibus (GEO) (http://www.ncbi.nlm.nih.gov/geo) under accession GSE158525.

## ACKNOWLEDGEMENTS

We thank members of the S.K. and D. L. laboratory for their helpful discussions.

## FUNDING

This work was supported by TJ Park Science Fellowship of the POSCO TJ Park Foundation (to S.E.P.) and the National Research Foundation of Korea [NRF-2018R1A5A1024261 to S.K. and D.L.].

## CONFLICT OF INTERESTS

The authors have no potential conflicts of interest to disclose.

---

**Figure 1—figure supplement 1.**
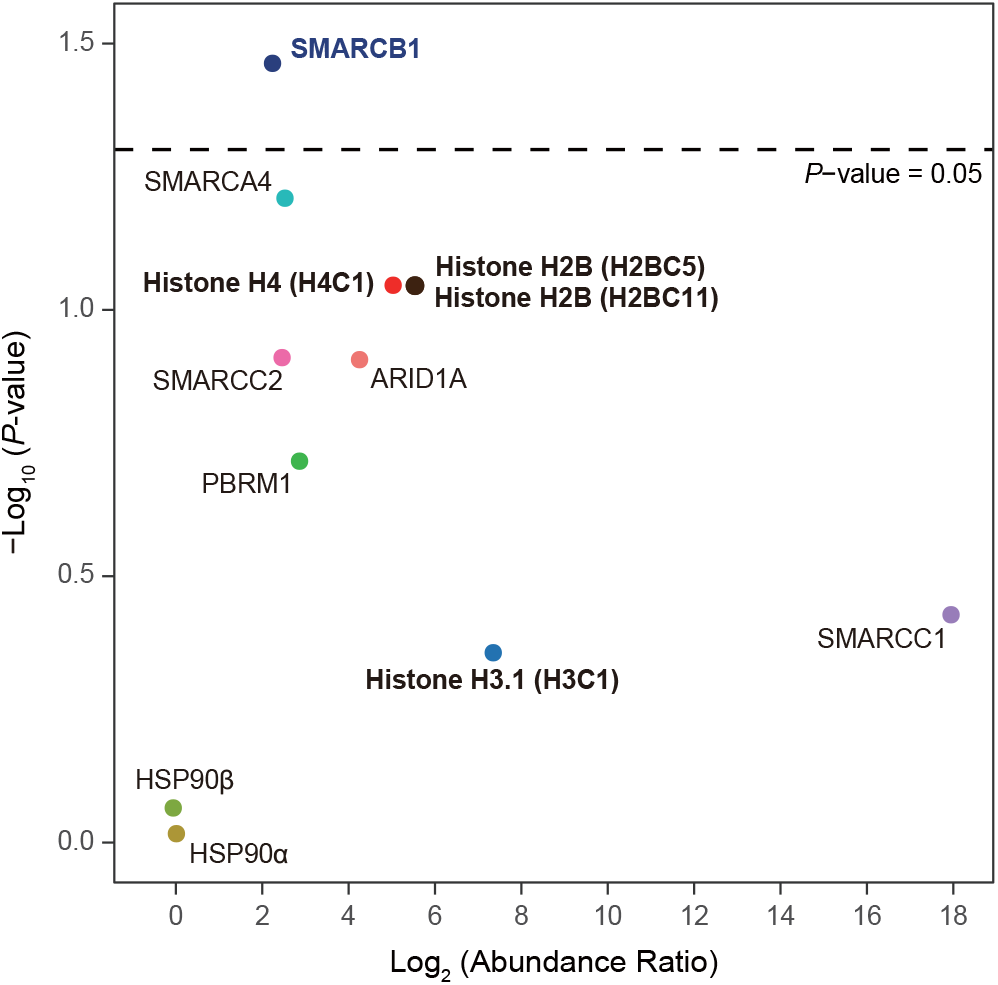
Various subunits of SWI/SNF complex and histones are IPMK-proximal/interacting proteins. A volcano plot showing the relative abundance and significance (*P*-value) of biotinylated proteins related to SWI/SNF complex, histones, and two negative controls (right). A dotted line within the volcano plot indicates the *P*-value = 0.05. The relative abundance (abundance ratio) was derived by comparing the fold enrichment of target proteins in IPMK-APEX2-expressed to APEX2-expressed HEK293 cells. *P*-value was calculated using Student’s t-test.

**Figure 2—figure supplement 1.**
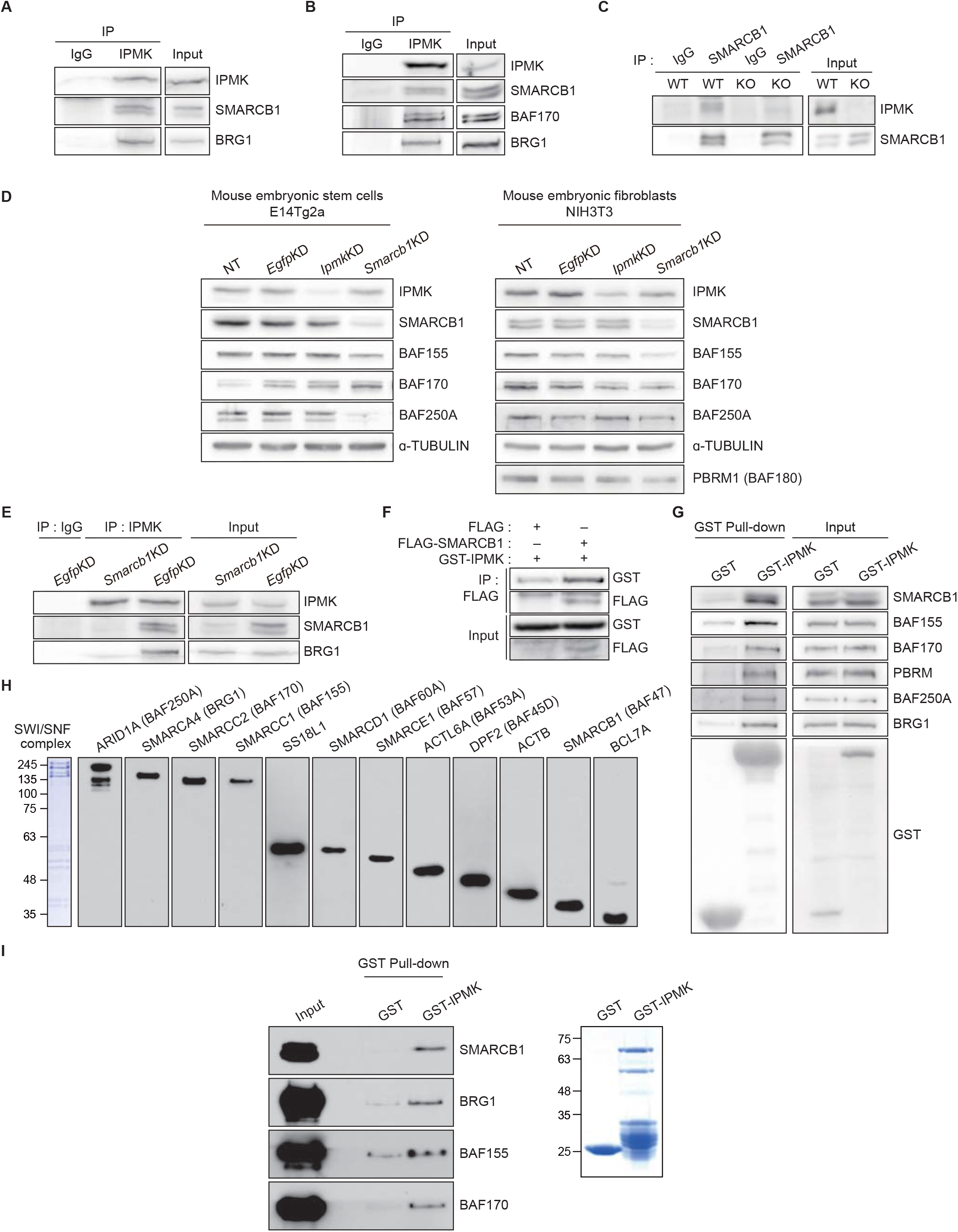
IPMK physically binds to SMARCB1 and other subunits of the SWI/SNF complex. (A) IPMK and IgG were immunoprecipitated from NIH3T3 cells and subjected to immunoblotting. (B) IPMK and IgG were immunoprecipitated from NIH3T3 cells in the presence of dithiobis (succinimidyl propionate) (DSP), an established crosslinker, and subjected to immunoblotting. (C) SMARCB1 and IgG were immunoprecipitated from wild-type (WT) and IPMK-depleted (KO) MEF cells and subjected to immunoblotting. (D) E14Tg2a cells and NIH3T3 cells were non-transfected (NT) or transfected with siRNA against *Egfp* (*Egfp*KD), *Ipmk* (*Ipmk*KD), and *Smarcb1* (*Smarcb1*KD), and then subjected to immunoblotting. (E) E14Tg2a cells were transfected with siRNA against *Egfp* (*Egfp*KD) and *Smarcb1* (*Smarcb1*KD), immunoprecipitated with IPMK and IgG, and subjected to immunoblotting. (F) HEK293T cells were co-transfected with GST-IPMK and FLAG-SMARCB1 or FLAG (a control vector), followed by FLAG immunoprecipitation and immunoblotting. (G) HEK293T cells were transfected with GST-IPMK or GST (a control vector), followed by GST pull-down and immunoblotting. (H) Coomassie blue staining (left) and immunoblots of native SWI/SNF complex purified from FLAG-DPF2 HEK293T cell line. (I) Native SWI/SNF complex purified from FLAG-DPF2 HEK293T cell line and purified GST-IPMK or GST were co-incubated, followed by GST pull-down and immunoblotting (left). Coomassie blue staining of purified GST and GST-IPMK proteins (right).

**Figure 3—figure supplement 1.**
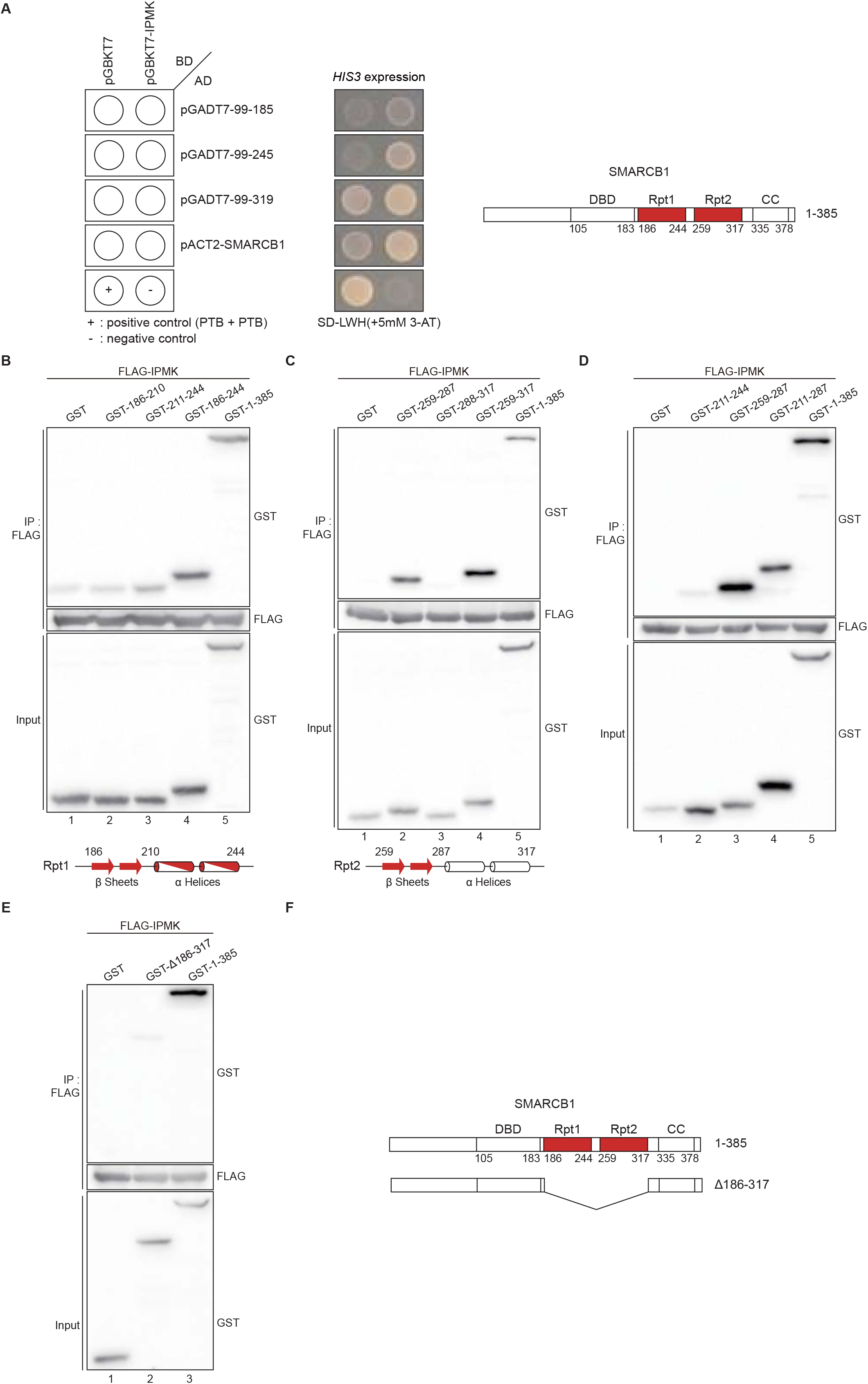
Domain mapping of the interaction between IPMK and SMARCB1. (A) IPMK and SMARCB1 domain interaction test in yeast strain AH109, containing the *HIS3* reporter gene. Yeast cells were co-transformed with either the GAL4-BD fusion plasmid pGBKT7 or pGBKT7-IPMK and the GAL4-AD fusion plasmid pGADT7 with SMARCB1 deletion constructs or pACT2-SMARCB1. IPMK interacts with 99-245 or 99-319 SMARCB1 deletion constructs, whereas IPMK does not interact with 99-185 SMARCB1 constructs. The yeast cells were spread on the selection medium lacking leucine and tryptophan (SD-LW) to select co-transformants of bait and prey vectors. Specific interactions between bait and prey proteins were monitored by cell growth on a selection medium lacking leucine, tryptophan, and histidine (SD-LWH). 3-AT (3-amino-1,2,4-triazole) was used to suppress leaky *HIS3* expression in transformants to obtain an accurate phenotype. A schematic diagram of the SMARCB1 domain map with the number of amino acid sequences is presented on the right. The IPMK-binding sites (Rpt1 and Rpt2) are highlighted in red. (B, C, and D) HEK293T cells were co-transfected with FLAG-IPMK and GST (a control vector) or GST-SMARCB1 Rpt1 fragments (B and D) and GST-MARCB1 Rpt2 fragments (C and D), followed by immunoprecipitation with FLAG antibody and subjected to immunoblotting. The specific IPMK-binding Rpt1 (B) and Rpt2 (C) domains are highlighted in red. Arrows and cylinders indicate β sheets and α helices, respectively. (E) HEK293T cells were co-transfected with FLAG-IPMK and GST (a control vector), GST-SMARCB1, or GST-SMARCB1 without Rpt1 and Rpt2 (GST-Δ186-371), followed by immunoprecipitation with FLAG antibody, and subjected to immunoblotting. (F) Schematic diagram of human SMARCB1 and SMARCB1 without Rpt1 and Rpt2 (Δ186-371). The IPMK-binding sites (Rpt1 and Rpt2) are highlighted in red with the number of amino acid sequences.

**Figure 4—figure supplement 1.**
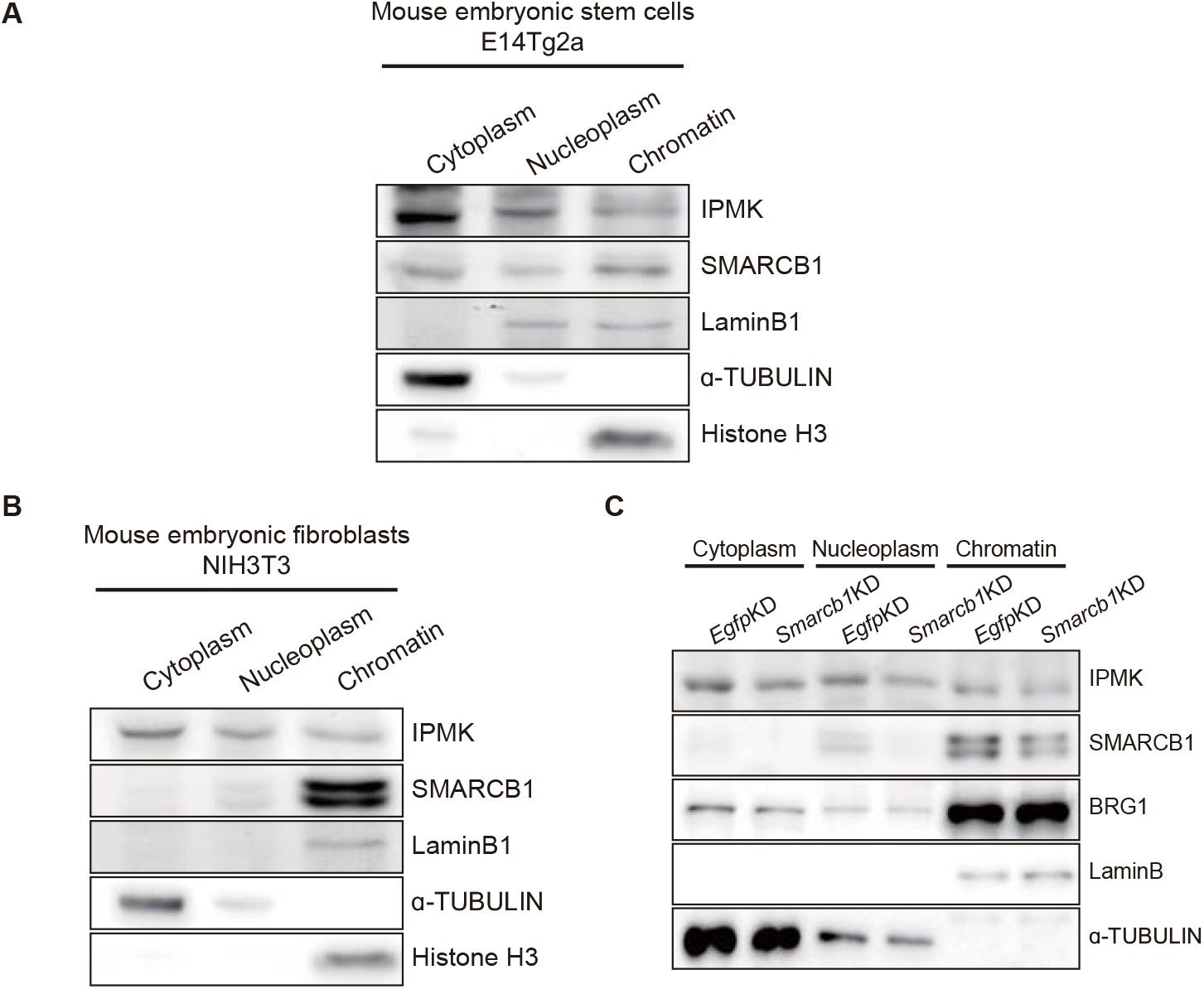
Chromatin fraction assay. (A) E14Tg2a cells were fractionated into the cytoplasm, nucleoplasm, and chromatin fractions. Immunoblotting with IPMK, SMARCB1, and fractionation markers was then performed. (B) NIH3T3 cells were fractionated into the cytoplasm, nucleoplasm, and chromatin fractions. Immunoblotting with IPMK, SMARCB1, and fractionation markers was then performed. (C) NIH3T3 cells were transfected with siRNA against *Egfp* (*Egfp*KD) and *Smarcb1* (*Smarcb1*KD) and then fractionated into the cytoplasm, nucleoplasm, and chromatin fractions. Immunoblotting with IPMK, SMARCB1, BRG1, and fractionation markers was then performed.

**Figure 5—figure supplement 1.**
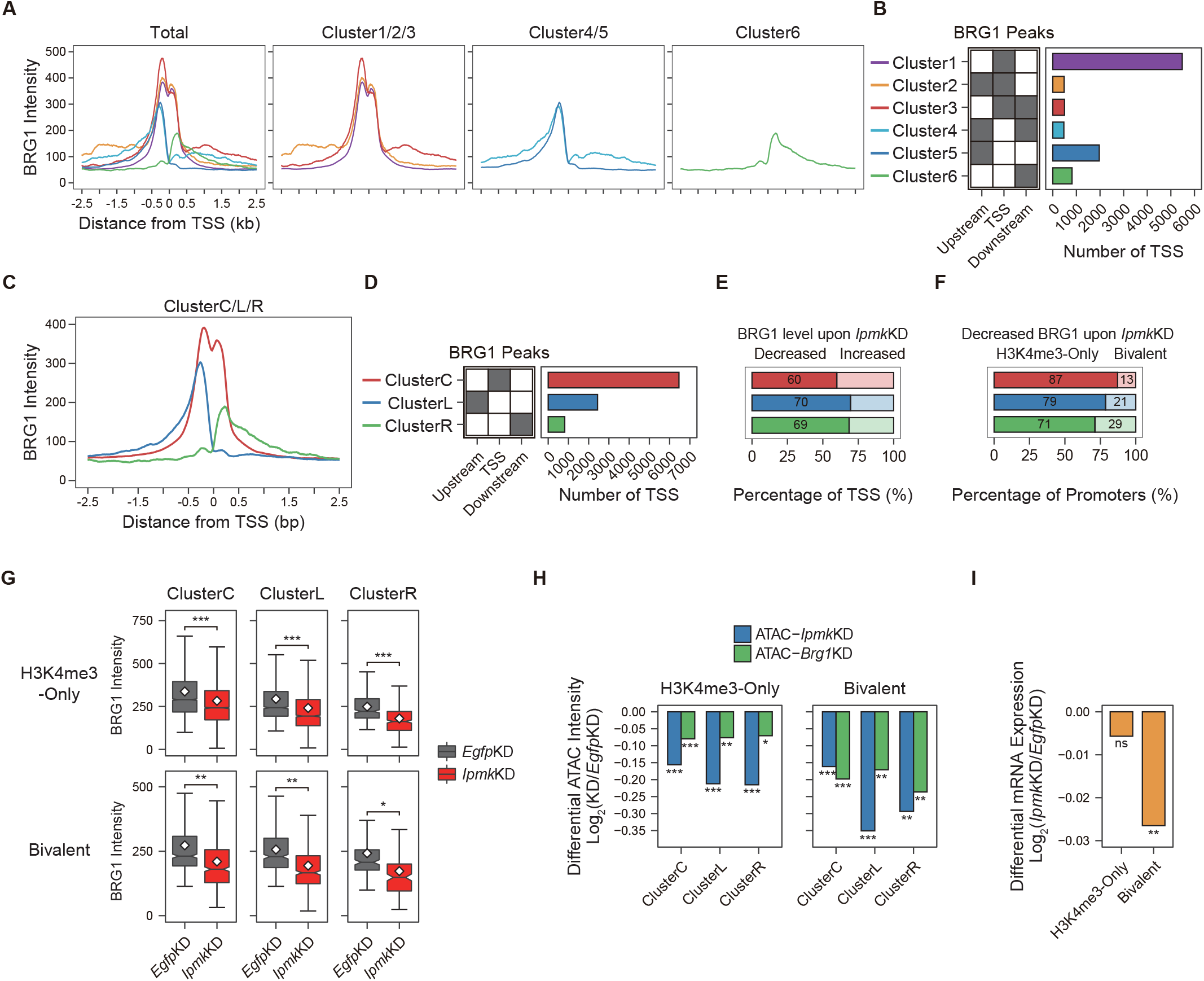
IPMK plays an important role in the maintenance of chromatin accessibility at promoter-TSS by regulating the BRG1 localization. (A) Line plots showing the average enrichments of BRG1 (*Egfp*KD cells) at TSS with six clusters. (B) A diagram displaying six clusters of TSS classified by the relative positions of BRG1 CUT&RUN peaks (*Egfp*KD cells) respective to TSS and up/downstream regions of TSS (left). Bar graphs showing the number of six TSS clusters (right). (C) Line plots showing the average enrichments of BRG1 (*Egfp*KD cells) at TSS with three clusters. ClusterC (red) contains cluster1-3, ClusterL (blue) contains cluster4-5, and ClusterR (green) resembles cluster6. (D) A diagram displaying three clusters of TSS classified by the relative positions of BRG1 CUT&RUN peaks (*Egfp*KD cells) respective to TSS and up/downstream regions of TSS (left). Bar graphs showing the number of three TSS clusters (right). (E) Bar graphs showing the percentage of three TSS clusters exhibiting decreased (left) or increased (right) BRG1 intensity upon *Ipmk*KD. (F) Bar graphs showing the percentage of three TSS clusters (TSS exhibiting decreased BRG1 intensity upon *Ipmk*KD) with H3K4me3-Only (left) and bivalent (right) promoters. (G) Box plots showing the BRG1 intensity at TSS with two promoter types (top: H3K4me3-Only promoters; bottom: bivalent promoters) and with three TSS clusters in *Egfp*KD (grey) and *Ipmk*KD (red) cells. *P*-values were derived using Wilcoxon signed rank test (*P < 1×10^-25^; **P < 1×10^-50^; ***P < 1×10^-100^). (H) Bar graphs showing the average of differential ATAC-seq intensity (Log_2_ KD/*Egfp*KD) upon *Ipmk*KD (blue) and *Brg1*KD (green) at three TSS clusters with H3K4me3-Only (left) and bivalent promoters (right). (I) Bar graphs showing the average of differential mRNA expression (Log_2_ *Ipmk*KD/*Egfp*KD) upon *Ipmk*KD at TSS (TSS exhibiting decreased BRG1 intensity upon *Ipmk*KD) with two promoter types. (H and I) *P*-values were derived using Wilcoxon signed rank test (*P < 0.01; **P < 1×10^-4^; ***P < 1×10^-10^; ns, not significant).

**Figure 6—figure supplement 1.**
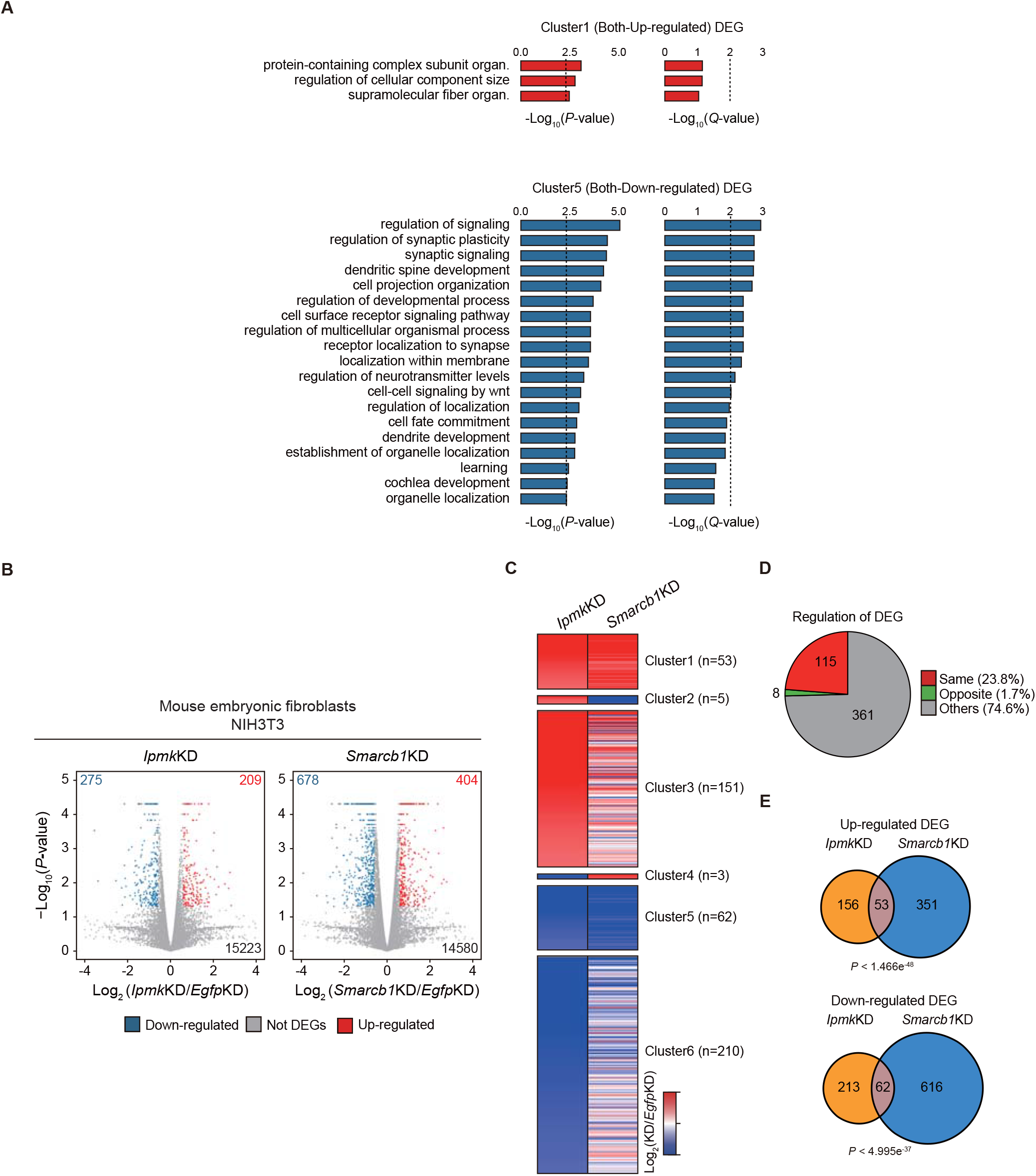
mRNA transcriptome indicates that IPMK and SMARCB1 regulate a common set of genes. (A) Bar graphs (left: *P*-value, right: *Q*-value) showing the gene ontology terms (biological process) of the Cluster1 (top) and Cluster5 (bottom) DEGs that satisfied the threshold (*P*-value ≤ 0.005). Organ. Denotes organization, and the dotted lines denote *P*-value ≤ 0.005 (left) and *Q*-value ≤ 0.01 (right). (B) Volcano plots showing the differentially expressed genes (DEGs) upon *Ipmk*KD (left) and *Smarcb1*KD (right) in NIH3T3 cells, identified based on mRNA-seq data. Red and blue dots indicate DEGs that were found to be significantly up-and down-regulated, respectively (*P*-value ≤ 0.05 and fold change ≥ 1.5). (C) Heatmaps representing differential gene expression (log_2_ (KD/*Egfp*KD)) of up-regulated (clusters1, 2, and 3) and down-regulated (clusters 4, 5, and 6) DEGs upon *Ipmk*KD. The heatmaps are classified into six clusters based on the differential gene expression of *Ipmk*KD and *Smarcb1*KD cells. (D) Pie chart showing the proportion of *Ipmk*KD-induced DEGs that are regulated in the same manner (red, clusters 1 and 5) or the opposite manner (green, clusters 2 and 4) upon *Ipmk*KD and *Smarcb1*KD. (E) Venn diagrams representing up-regulated DEGs (top) and down-regulated DEGs (bottom) in *Ipmk*KD (orange) and *Smarcb1*KD (blue) cells. The *P*-values indicate the significance of the overlap between the two groups.

**Figure 7—figure supplement 1.**
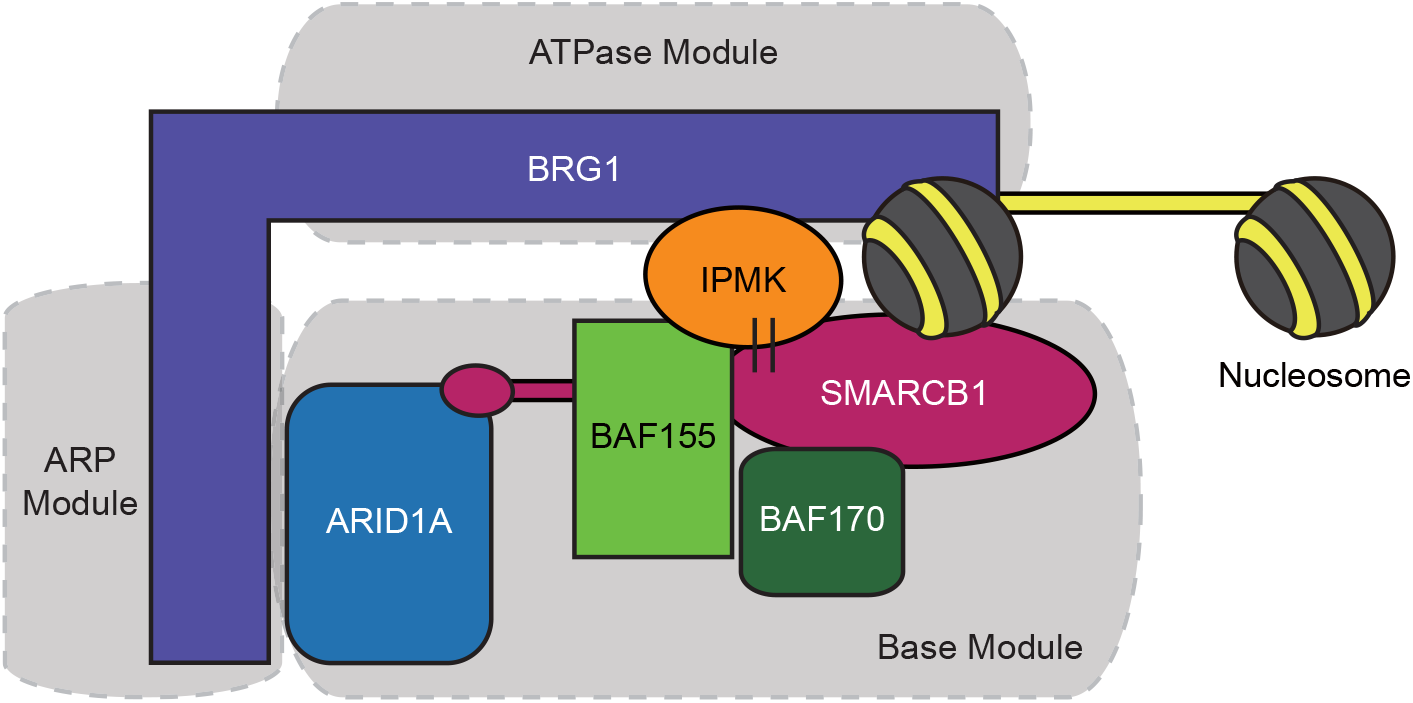
Proposed model depicting the physical interaction of IPMK and nucleosome-bound SWI/SNF complex. An additional model showing our speculation on position of IPMK within the SWI/SNF complex in the presence of nucleosomes.

**Supplementary Table 1.**
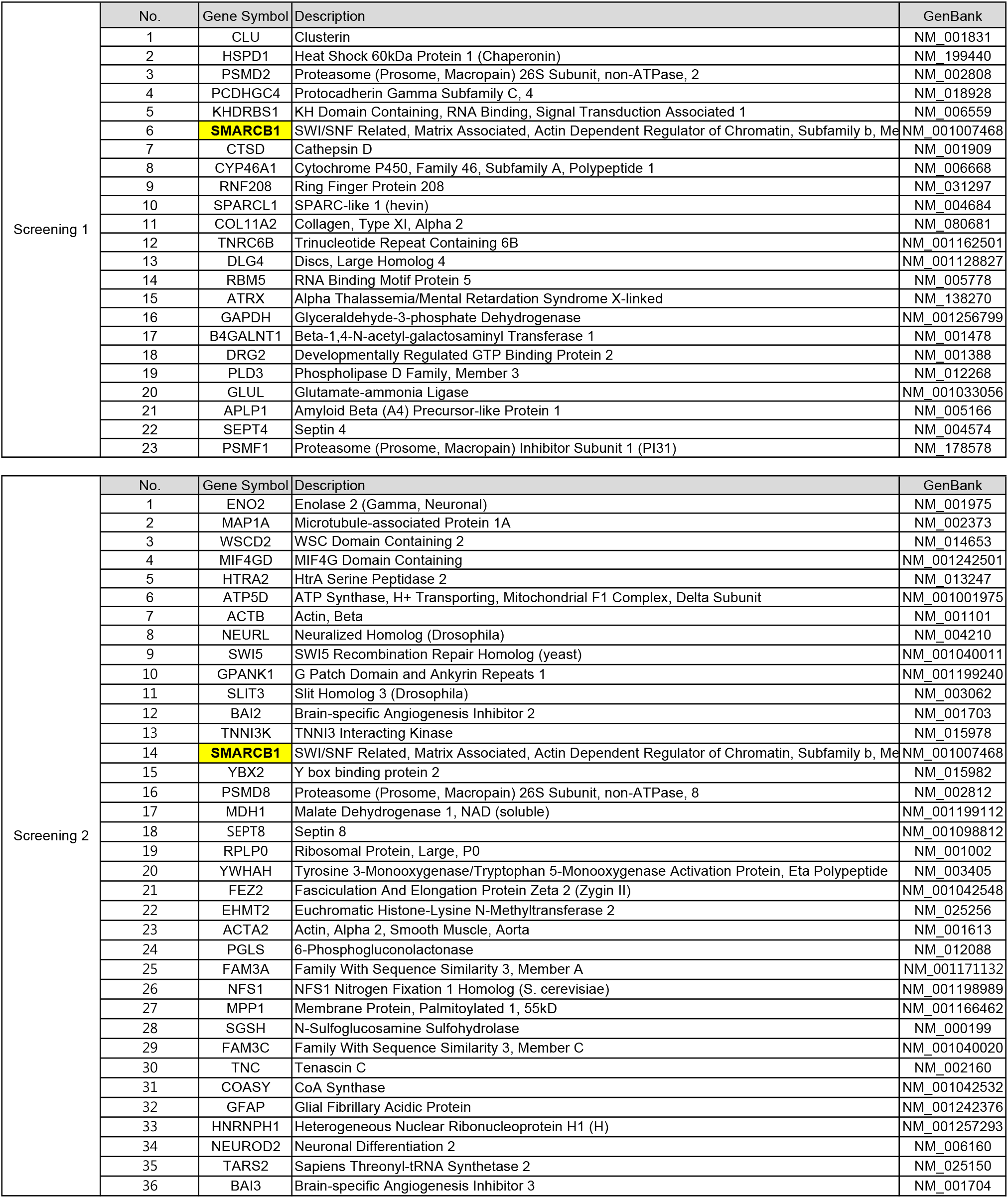
Yeast two-hybrid screening assay using IPMK as bait.

**Supplementary Table 2.** Enriched protein complex-based sets from APEX2-mediated proximity labeling.

| Complex name | Set size | Candidates contained | q-value |
| --- | --- | --- | --- |
| BRG1-SIN3A-HDAC containing SWI/SNF remodeling complex I | 11 | 7 (63.6%) | 4.83E-07 |
| BRG1-SIN3A complex | 14 | 7 (50.0%) | 4.00E-06 |
| Brg1-associated complex II | 7 | 5 (71.4%) | 1.60E-05 |
| BRG1-associated complex | 9 | 5 (55.6%) | 6.98E-05 |
| BAF complex | 9 | 5 (55.6%) | 6.98E-05 |
| PBAF complex (Polybromo- and BAF containing complex) | 9 | 5 (55.6%) | 6.98E-05 |
| BRM-SIN3A complex | 15 | 6 (40.0%) | 8.10E-05 |
| EBAFa complex | 10 | 5 (50.0%) | 0.000104312 |
| BAF complex | 13 | 5 (38.5%) | 0.000329586 |
| Brg1-based SWI/SNF complex | 4 | 3 (75.0%) | 0.000720336 |
| EBAFb complex | 10 | 4 (40.0%) | 0.001110216 |
| BRM-associated complex | 10 | 4 (40.0%) | 0.001110216 |

**Supplementary Table 3.**
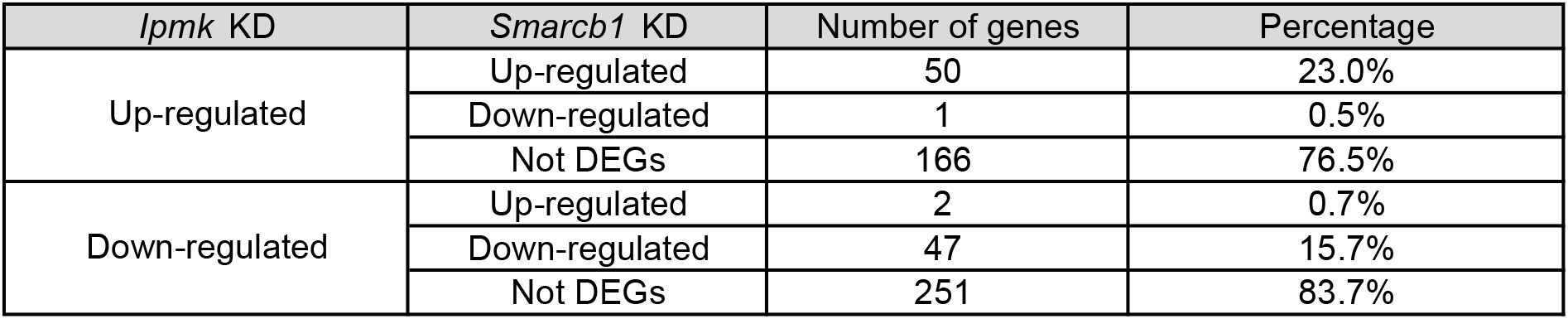
Clustering analysis of DEG in mouse embryonic stem cells.

**Supplementary Table 4.**
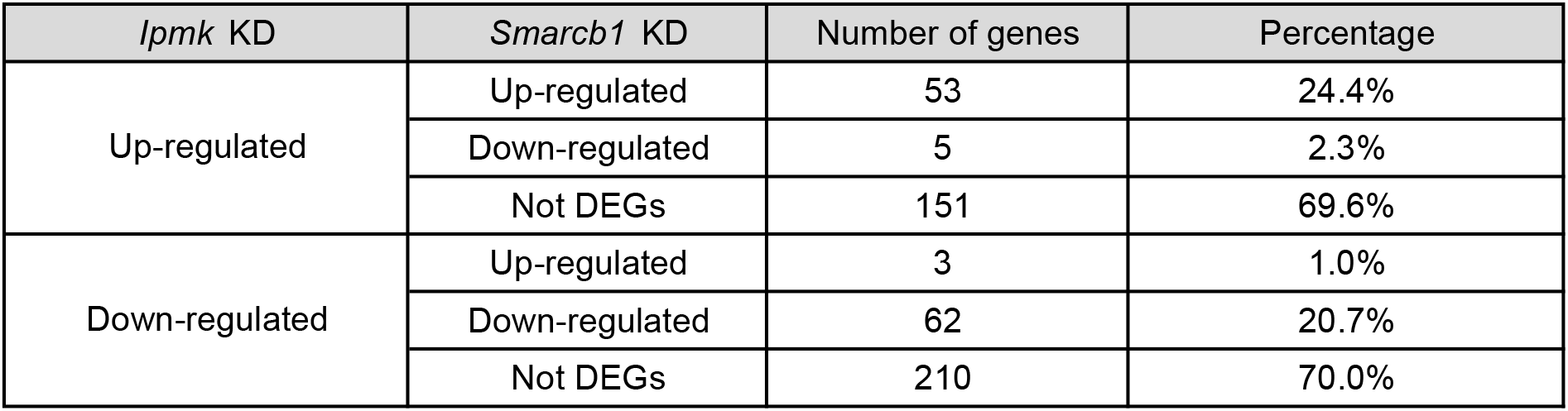
Clustering analysis of DEG in mouse fibroblasts.

## Notes

### Competing Interest Statement

The authors have declared no competing interest.

